# Single-nuclei transcriptomics revealed auxin-driven mechanisms of wood plasticity and severe drought tolerance in poplar

**DOI:** 10.1101/2025.01.08.631930

**Authors:** Daniela Gómez-Soto, Wendell J. Pereira, Alejandro Piedrabuena-Díaz, Christopher Dervinis, Matias Kirst, Isabel Allona, Mariano Perales, Daniel Conde

## Abstract

Drought significantly affects forests and woody crops by limiting their growth, increasing their susceptibility to diseases, and reducing productivity. Wood anatomical plasticity is a crucial adaptive mechanism that enables trees to cope with fluctuations in water availability. During severe drought, trees develop more and narrower vessels, enhancing hydraulic safety and reducing the risk of embolism. However, the molecular regulation of vessel formation is still not well understood. Using single-nucleus transcriptomics, we generated a cell type-specific gene expression map of the mature poplar stem under well-watered and drought conditions. Our findings revealed extensive gene expression reprogramming in xylem-forming cells, with changes in auxin homeostasis identified as a key mechanism for anatomical adaptation. Specifically, we showed that poplar *WAT1*-like genes control vessel spatial patterning. Additionally, the downregulation of *WAT1*-like gene expression in the dividing cells of the vascular cambium and the upregulation of *MP*-like gene in cells undergoing early vessel differentiation facilitate the formation of secondary xylem with narrower and more numerous vessels. Furthermore, the *wat2* mutant exhibited greater drought tolerance than wild-type trees, underscoring its potential for developing drought-resilient tree varieties. These insights enhance our understanding of xylem plasticity and provide valuable targets for improving drought tolerance in woody plants.

## BACKGROUND

Global climate change, driven by human emissions of greenhouse gases, is expected to have critical implications for tree survival^1,2^. The elevated temperature of the Earth’s surface results in a higher evaporation rate and increased vapor circulation in the atmosphere, disrupting the temporal and spatial distribution of precipitation. The water cycle intensification is projected to lead to more frequent and severe climatic events, including extreme droughts^3^. Access to water impacts tree phenology and growth^4^ and, consequently, the productivity of trees and forests^5^.

In trees and other vascular plants, drought episodes can irreversibly impact the functionality of the vascular system^6^. In trees, the shoot apical meristem (SAM) activity establishes the primary vascular meristem or procambium. Procambial cell divisions give rise to the primary xylem and phloem, forming the plant’s primary vascular system^7^. In species with secondary growth (gymnosperms and many dicotyledons), the secondary vascular meristem, the vascular cambium, is derived from subsets of procambial cells. Secondary growth involves increasing vascular tissues by forming wood (secondary xylem) and secondary phloem^8^. The secondary xylem forms the vast majority of tree biomass and vasculature, and it is responsible for providing structural support and water supply from the root system to the shoot. In perennials, wood is formed by an intricate network of vessels, fibers, and parenchymatic cells. The xylem comprises mature, dead tracheary elements that connect to form long strands used to conduct water and minerals from the roots to the leaves. The two fundamental types of xylem conduits are tracheids (typical to gymnosperms) and vessels (in angiosperms) built of vessel elements. Vessels and fibers share similar developmental stages during their differentiation^9^ and few genes specific to each cell type have been characterized. Several VASCULAR-RELATED NAC-DOMAIN genes regulate secondary cell wall formation of vessels (SCW), while the *ACAULIS* 5 (*ACL5*) thermospermine synthase regulates vessel programmed cell death (PCD)^10^. The gene *ENLARGED VESSEL ELEMENT* contributes to vessel dimension in poplar^11^.

In trees, including poplar, long-term drought modifies the pattern of vessel formation^12,13^. Drought induces the formation of extra vessel cells with reduced cellular diameter, increasing hydraulic safety and minimizing the risk of embolism and die-back^14–17^. Despite their tremendous importance in tree adaptation, vessel diameter and frequency regulation are among the most poorly understood aspects of xylem differentiation. Understanding the specific pathways that regulate vessel element number and size is crucial to understanding how transcriptional changes during drought translate into compensating features of wood anatomy. This knowledge would allow altering these pathways to engineer more resilient tree species.

To gain insights into the molecular drivers of wood plasticity under severe drought, we applied single nucleus transcriptomics (snRNA-seq). First, we generated a cell type-specific gene expression map of the mature poplar stem under both drought and well-watered (control) conditions. Next, we contrasted the developmental trajectories of secondary xylem formation and identified the gene expression reprogramming that occurs under severe drought in each cell type involved in wood formation. This work establishes the first comparison of secondary xylem development at the single-cell resolution between well-watered and severe drought-treated plants in hybrid poplar.

## RESULTS

### Severe drought causes a reconfiguration of xylem patterning in the hybrid poplar stem

We compared the secondary xylem formed during a 20-day drought period in which the soil was maintained at 15% of its maximum water-holding capacity, also called field capacity (FC), to xylem developed when plants were well-watered (control). We observed a reconfiguration of the xylem cellular pattern in the newly formed wood established under drought compared to well-watered conditions (Fig. 1A, B and Supplementary Fig. 1). We counted the number of cells of each type formed in secondary xylem (internode 15) developed under both growth conditions, before and after the drought treatment. The vessel ratio, defined as the proportion of vessel to fiber cells, ranged from 5% to 7% (Supplementary Fig. 1) in the wood formed during the well-watered period, with an average of 6% (Fig. 1C). The vessel ratio significantly increased in the wood formed under drought in all individuals, ranging from 13% to 22% (Supplementary Fig. 1), with an average of 17% (Fig. 1C). Furthermore, vessel lumina was reduced considerably in these plants (Fig. 1A). Our observations indicate that these poplar trees generate wood with smaller and more frequent vessels in response to drought.

**Fig. 1.**
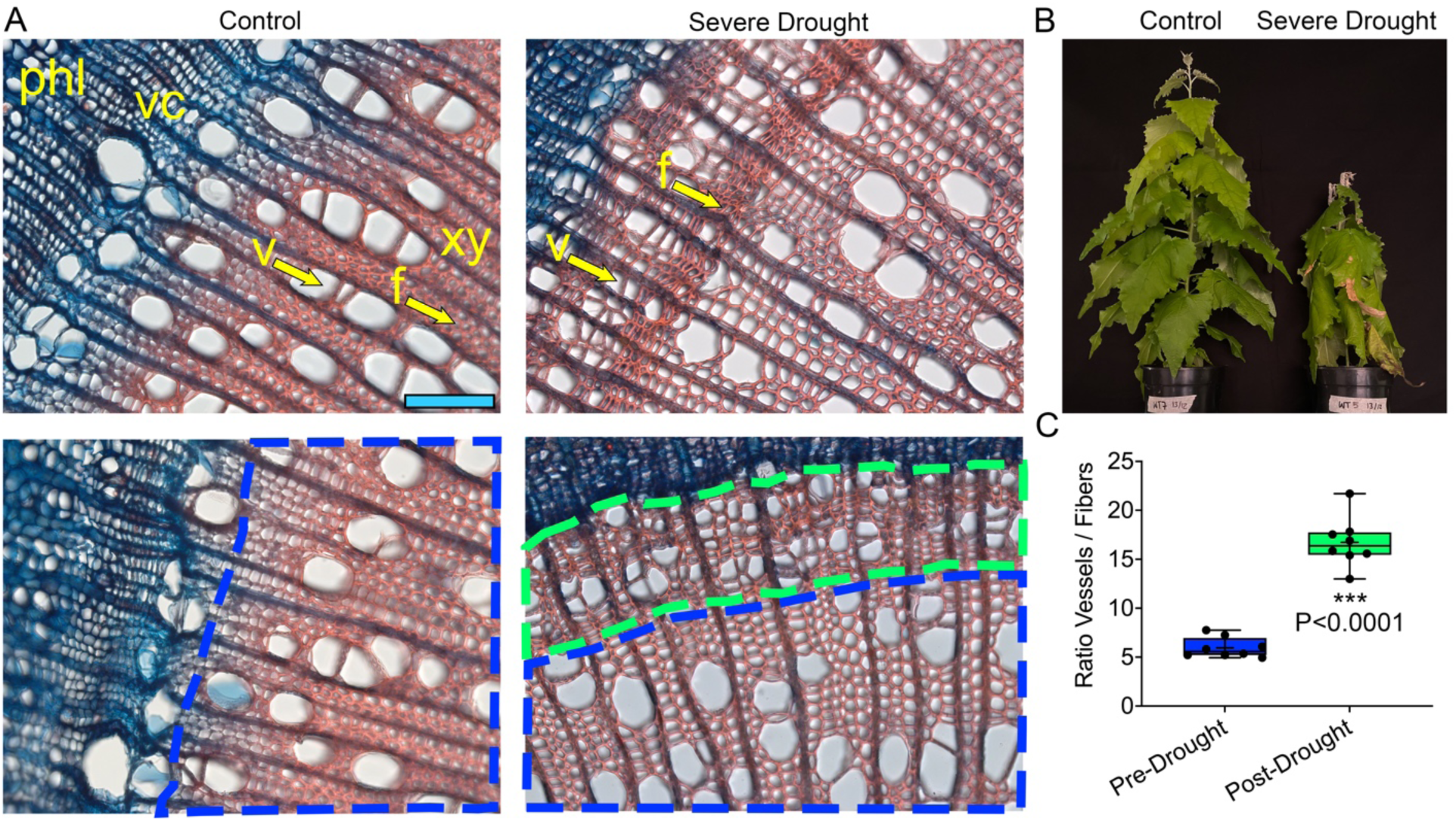
Characterization of secondary xylem formation under severe drought. **A** Stem cross-sections of internode 15 of WT trees growing under well-watered (control) and drought conditions, stained with safranine (secondary cell wall) and alcian blue (primary cell wall). The blue dashed line frames the secondary xylem formed during the period without water deficiency, while the xylem formed under the drought period is framed by the green dashed line. Phloem (phl), vascular cambium (vc), xylem (xy), vessels (v), and fibers (f) are labeled. **B** Representative trees of the individuals subjected to the well-watered and severe drought treatments after 14 days. **C** Under severe drought, the secondary xylem significantly increases the ratio of vessels/fibers. For each box-and-whisker plot, the center black line and the “+” represent the median and the mean, respectively; the box extends from the 25th to 75th percentiles; the whiskers are the maximum and the minimum values (*n*=8 trees). Student’s t-test determined statistical differences. Scale bar 100 μm in A.

### A cell type-specific transcriptome of hybrid poplar mature stem under well-watered and drought conditions

To identify molecular drivers of the xylem response to drought, we carried out snRNA-seq of poplar stems sampled from plants grown under well-watered and drought conditions (14 days with the FC at 15%). Four snRNA-seq libraries, two per condition, were generated. For each library, nuclei were isolated from the internode 15 from eight-week-old hybrid poplars. After filtering for good-quality nuclei, a comparison of each library’s average gene expression showed a high correlation between biological replicates while highlighting transcriptome changes between well-watered and drought treatments (Supplementary Fig. 2). Using Asc-Seurat^18^, data from each treatment and replicate were integrated to obtain a combined dataset containing 14,004 nuclei, with 7,089 nuclei coming from the well-watered trees and 6,915 from drought-treated trees (Fig. 2A). On average, 981 and 784 expressed genes were identified per nucleus for the well-watered and drought treated samples, respectively. Overall, the expression of 25,109 genes was detected. Before characterizing gene expression reprogramming under severe drought, we annotated the clusters obtained by unsupervised graph-based clustering, implemented in Asc-Seurat (Fig. 2A). We based the cluster annotation on the expression patterns of previously characterized cell-type-specific genes (Fig. 2B and Supplementary Table 1). Transcripts of the cambial stem cell regulators *AINTEGUMENTA* (*ANT*) and *ANT*-like 5^19,20^, were enriched in cluster 10 (Fig. 2B). Cambium cells differentiating into xylem were identified based on the expression of *WOX4* (*WUSCHEL-RELATED HOMEOBOX 4*)^21^ and *PXY* (*PHLOEM INTERCALATED WITH XYLEM*)^22,23^, enriched in clusters 0, 3, 7 and 13 (Fig. 2B). Additional evidence suggesting that clusters 3 and 7 contain cells undergoing xylem differentiation was inferred from the expression of *ACAULIS 5* (*ACL5*) and *PIN-FORMED 6* (*PIN6*)^24,25^. The expression of the transcriptional regulator of secondary cell wall (SCW) formation, *VASCULAR RELATED NAC-DOMAIN PROTEIN 1* (*VND1*), was specifically identified in cluster 6^26^. Other genes involved in SCW biosynthesis were highly expressed in clusters 6, 19 and 22 (Fig. 2B), including *KNOTTED ARABIDOPSIS THALIANA 7* (*KNAT7*), *CELLULOSE SYNTHASE 8* (*CESA8*), *LACCASE 4* (*LAC4*) and *LACCASE 17* (*LAC17*), which act redundantly during vessel element and fiber lignification^27^. The expression of genes involved in lignin biosynthesis, such as *4-COURMARATE:COA LIGASE* (*4CL*), were also detected in those clusters^28^. Poplar *XYLEM CYSTEINE PEPTIDASE 2* (*XCP2*), a marker for xylogenesis that controls vessel autolysis during programmed cell death (PCD), was also mainly expressed in cluster 22^29^. This observation suggests that xylem vessel elements and fibers undergoing secondary cell wall formation and programmed cell death in the secondary xylem are represented by nuclei in clusters 6, 19, and 22 (Fig. 2B).

**Fig. 2.**
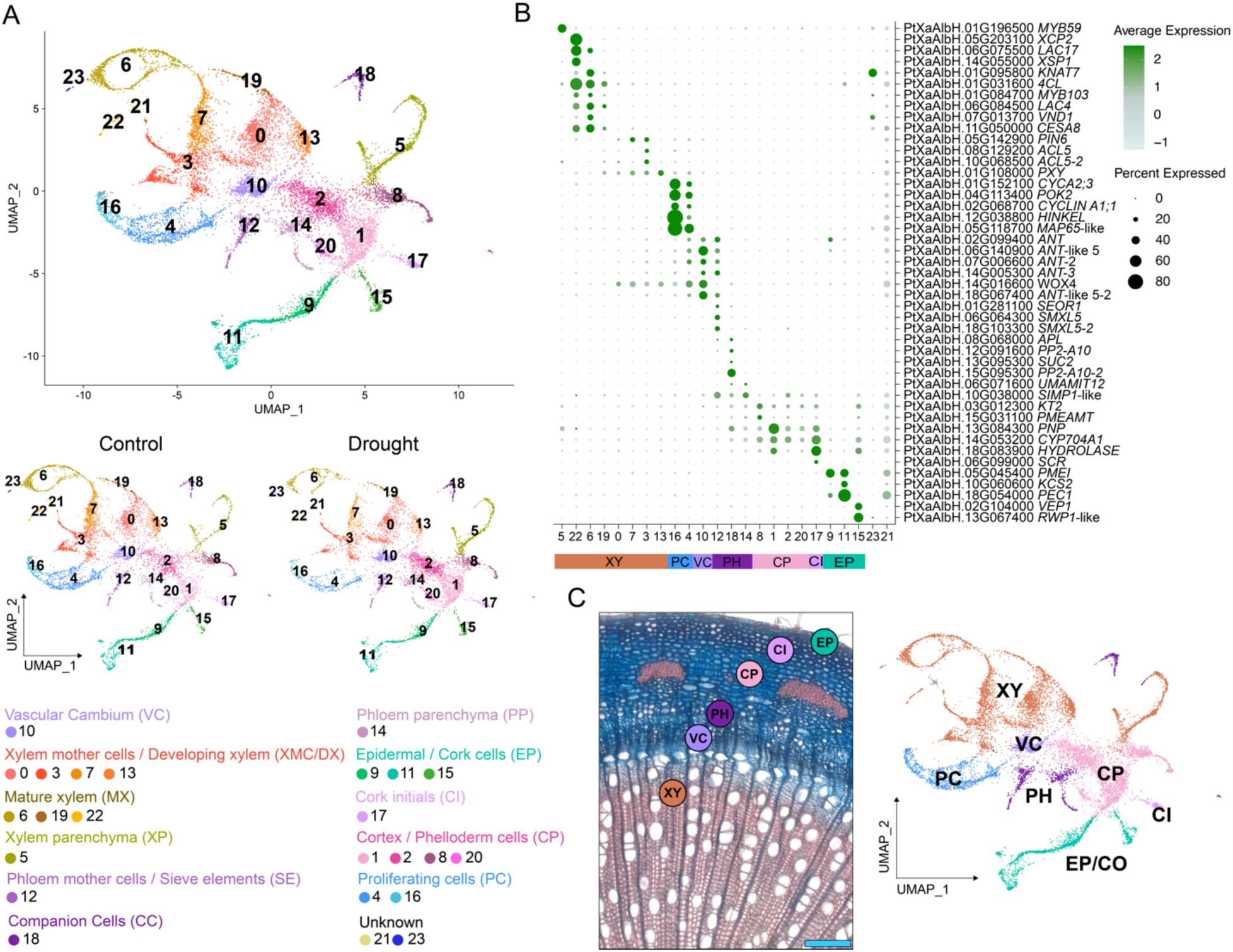
Cell type-specific transcriptome in the poplar stem under well-watered and drought conditions. **A** Visualization of the 24 cell clusters using UMAP after integrating the snRNA-seq data. Dots, individual cells; 7,089 cells for the well-watered (control) and 6,915 for the severe drought; color, cell clusters. **B** Expression pattern of previously characterized cell type marker genes in the hybrid poplar stem snRNA-seq dataset. Dot diameter is the proportion of cluster cells expressing the gene. **C** A stem-cross section of a hybrid poplar stem stained with safranin and alcian blue shows the spatial location of the seven annotated cell types and their visualization in the dataset using UMAP. Scale bar 250 μm in C.

We also captured the phloem mother cells (PMC, cluster 12) and their derivatives: companion cells (CC, cluster 18), sieve elements (SE, cluster 12), and phloem parenchyma (PP, cluster 14). Cluster 12 contains the PMC, expressing genes that promote phloem initials from cambial cells, such as *SMAX2 1*-LIKE 5^30^. Expression of SE markers such as *SIEVE ELEMENT OCCLUSION-RELATED 1* (*SEOR1*)^31^ was also enriched in cluster 12. Several genes associated with the SE-CC complex, including members of the *PHLOEM PROTEIN 2* family^32^ (*PP2-A1* and *PP2-A10*) and *SUCROSE-PROTON SYMPORTER 2* (*SUC2/SUT1*) involved in the uptake of sucrose in CC were primarily expressed in cluster 18. To annotate clusters representing PP cells, we evaluated the expression of *SALT INDUCED MALECTIN-LIKE DOMAIN-CONTAINING PROTEIN1* (*SIMP1*) and *UMAMIT 12*^33^.

During stem growth and maturation, the epidermis is replaced by the periderm, which is composed of a secondary meristem. Two tissues develop from this meristem: phelloderm, deposited internally, and phellem or cork tissue, found externally. Cork– and epidermis-specific markers analysis indicated that clusters 9, 11, and 15 contain these cells. The expression of cork markers *PECTIN METHYLESTERASE INHIBITOR* (*PMEI*) and *PERMEABLE CUTICLE 1* (*PEC1*) are enriched in clusters 9 and 11 (Fig. 2B)^34^. Epidermis-specific genes, such as *3-KETOACYL-COA SYNTHASE 2* (*KCS2*), were enriched in cluster 11^35^. Also, the expression of the poplar epidermal markers *VEIN PATTERNING 1* (*VEP1*) and *REDUCED LEVELS OF WALL-BOUND PHENOLICS 1*-like (*RWP1*-like) are enriched in cluster 15 (Fig. 2B)^34^. We found *SCARECROW* (*SCR*) expression to be specific to cluster 17. This gene controls tissue patterning by regulating the formative asymmetric division and, in poplar, is expressed at the cork initials^36^. To identify clusters of cells corresponding to the phelloderm/cortex (collenchyma), we explored the expression of markers identified in the previously published single-cell expression database of poplar stem^34^. We found markers of phelloderm mainly expressed in clusters 1 and 8 and with less expression level in clusters 2 and 20 (Fig. 2B and Supplementary Table 1). We hypothesized that these clusters contain the cortex/phelloderm cells (Fig. 2C).

### Cell type-specific gene expression reorganization during secondary xylem formation in response to severe drought

To investigate the gene regulatory network reprogramming that governs wood plasticity and confers adaptation to drought, we analyzed the cells involved in secondary xylem formation separately. The secondary xylem arises from the cell division of vascular cambium cells, which create the daughter cells that undergo cell differentiation. In the case of fibers and vessels, this comprises a synchronized and sequential program of cell expansion, SCW formation/lignification, and PCD. From the initial dataset (Fig. 2A) we reclustered the nuclei annotated as Vascular Cambium (VC) (cluster 10), Proliferating Cells (PC) (clusters 4 and 16), Xylem Mother Cells / Developing Xylem (XMC/DX) (clusters 0, 3, 7 and 13), Mature Xylem (MX) (clusters 6, 19 and 22), and Xylem Parenchyma (XP) (cluster 5) (Fig. 3A and Supplementary Fig. 3). Exploration of well-known cell type markers after reclustering enabled the identification of cells at different stages of secondary xylem formation, from vascular cambium and dividing cells to cells undergoing SCW formation and PCD (Supplementary Fig. 4). This xylem development reclustering data (XDR) resulted in seven clusters annotated as VC (cluster 5), PC (cluster 2), XMC / DX (clusters 0 and 1), MX (clusters 3 (SCW) and 6 (SCW/PCD)), and XP (cluster 4) (Fig. 3A and Supplementary Fig. 3).

**Fig. 3.**
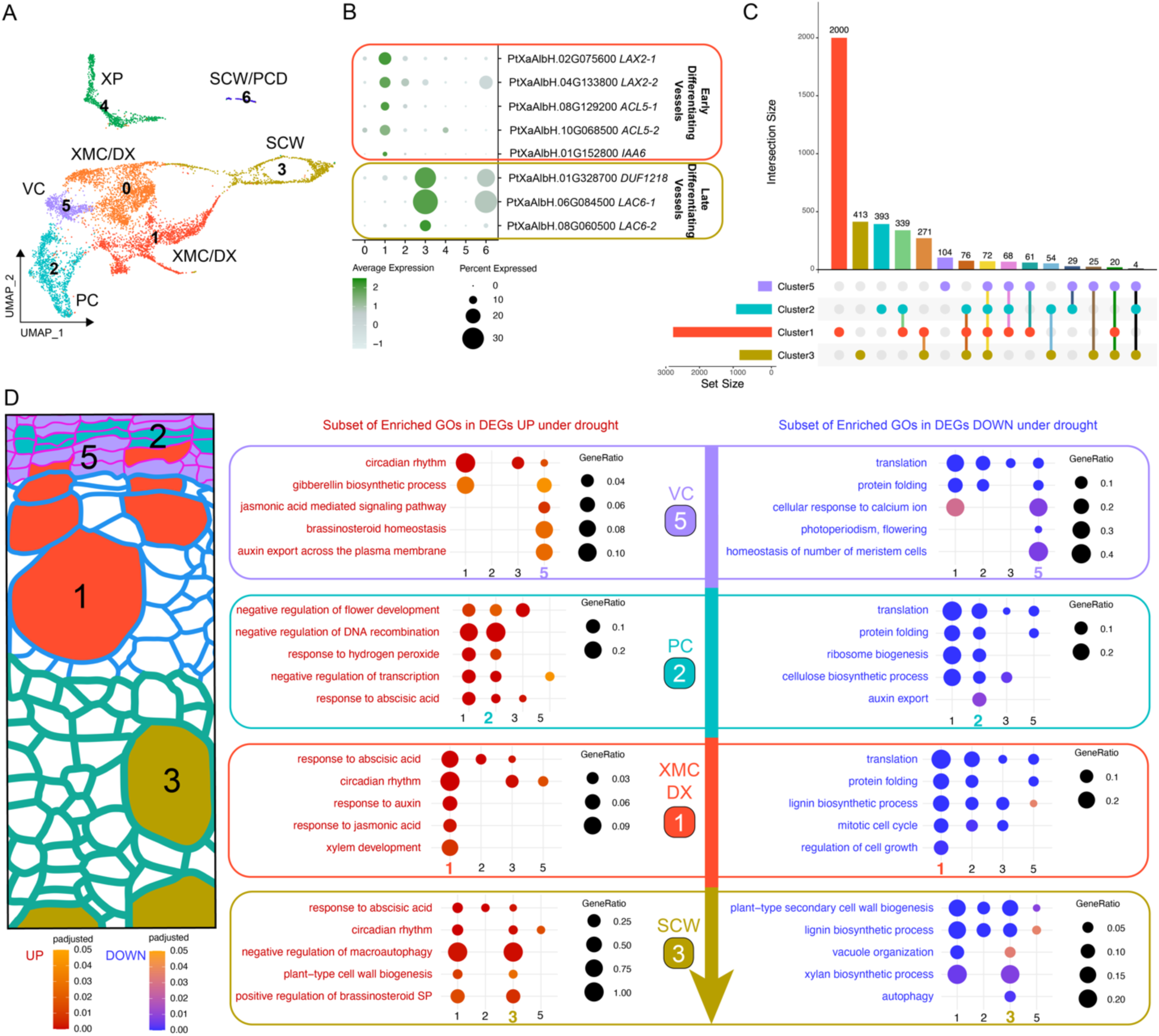
Cell-type specific reprogramming under severe drought. **A** Reclustering of the cells involved in secondary xylem formation: Vascular Cambium (VC), Proliferating Cells (PC), Xylem Mother Cells/Developing Xylem (XMC/DX), Xylem Parenchyma (XP), Secondary Cell Wall (SCW) and Programmed Cell Death (PCD). **B** Exploration of the expression of marker genes for Early Developing Vessels and Late Developing Vessel highlighted their enrichment in cluster 1 and cluster 3, respectively. **C** The UpSet plot displays the distribution and intersection of DEGs identified at clusters involved in vessel differentiation (clusters 1, 2, 3, and 5) under severe drought, respect the control. The horizontal bars indicate the total number of DEGs for each cluster. The vertical bars represent the number of DEGs specific across intersecting sets, with the dots connected by lines below identifying the specific intersections represented in each vertical bar. **D** Depiction showing the cells involved in the vessel differentiation in a stem cross-section during secondary xylem formation, from VC to SCW formation, and a subset of enriched Biological Processes in each list of DEGs for cluster 5 (VC), cluster 2 (PC), cluster 1 (XMC enriched in Early Differentiating Vessels) and cluster 3 (SCW).

Recently, a scRNA-seq study of the *Arabidopsis* root during secondary growth identified and characterized markers specific to xylem fibers and vessels^37^. We examined the expression of hybrid poplar homologs of the genes that served as markers for early differentiating vessels in *Arabidopsis*, and found their expression enriched in cluster 1 (Fig. 3B). Additionally, markers of late differentiating vessels from *Arabidopsis* showed enriched expression in clusters 3 and 6 (Fig. 3B), corresponding to cells undergoing SCW formation and PCD. These findings suggest that among clusters containing XMC/DX (clusters 0 and 1), cluster 1 is primarily enriched for cells undergoing vessel element differentiation. Furthermore, as vessel differentiation proceeds, the SCW formation induces substantial shifts in the gene expression network. Consequently, xylem cells undergoing SCW formation, including vessels and fibers, are localized within clusters 3 and 6 (Fig. 3A).

The developmental program that specifically drives vessel formation, rather than fiber differentiation, remains poorly understood. *ACL5* is one of the few genes specifically expressed in cells undergoing early vessel differentiation in the *Arabidopsis* root^24,37^. The hybrid poplar genome contains three *ACL5* homologs (*ACL5*-like 1; PtXaAlbH.06G184800, *ACL5*-like 2; PtXaAlbH.08G129200, and *ACL5*-like 3; PtXaAlbH.10G068500), located on chromosomes 6, 8, and 10, respectively. Under well-watered conditions, these genes are primarily expressed in PC and a specific region of cluster 1 (XMC/DX) in the XDR data (Fig. 4A and Supplementary Fig. 5), with PtXaAlbH.06G184800 also showing notable expression in XP (Supplementary Fig. 5). Under drought conditions, PtXaAlbH.08G129200 expression remains mainly localized to the same cells in cluster 1 (Fig. 4A), while PtXaAlbH.06G184800 and PtXaAlbH.10G068500 exhibit increased expression in XP and at a specific region of cluster 0 (Supplementary Fig. 5). Due to its higher specificity, we generated stable transgenic lines expressing an *H2B-YFP* reporter driven by the PtXaAlbH.08G129200 promoter. Fluorescence in the stem cross-section was localized to specific cells in the cambial, developing xylem regions, and developing vessels (Fig. 4B). These findings suggest that similar to *Arabidopsis*, *ACL5*-like 2 is predominantly expressed in cells at the early stages of vessel element differentiation. This supports our hypothesis that cluster 1 is enriched in vessel-developing cells.

**Fig. 4.**
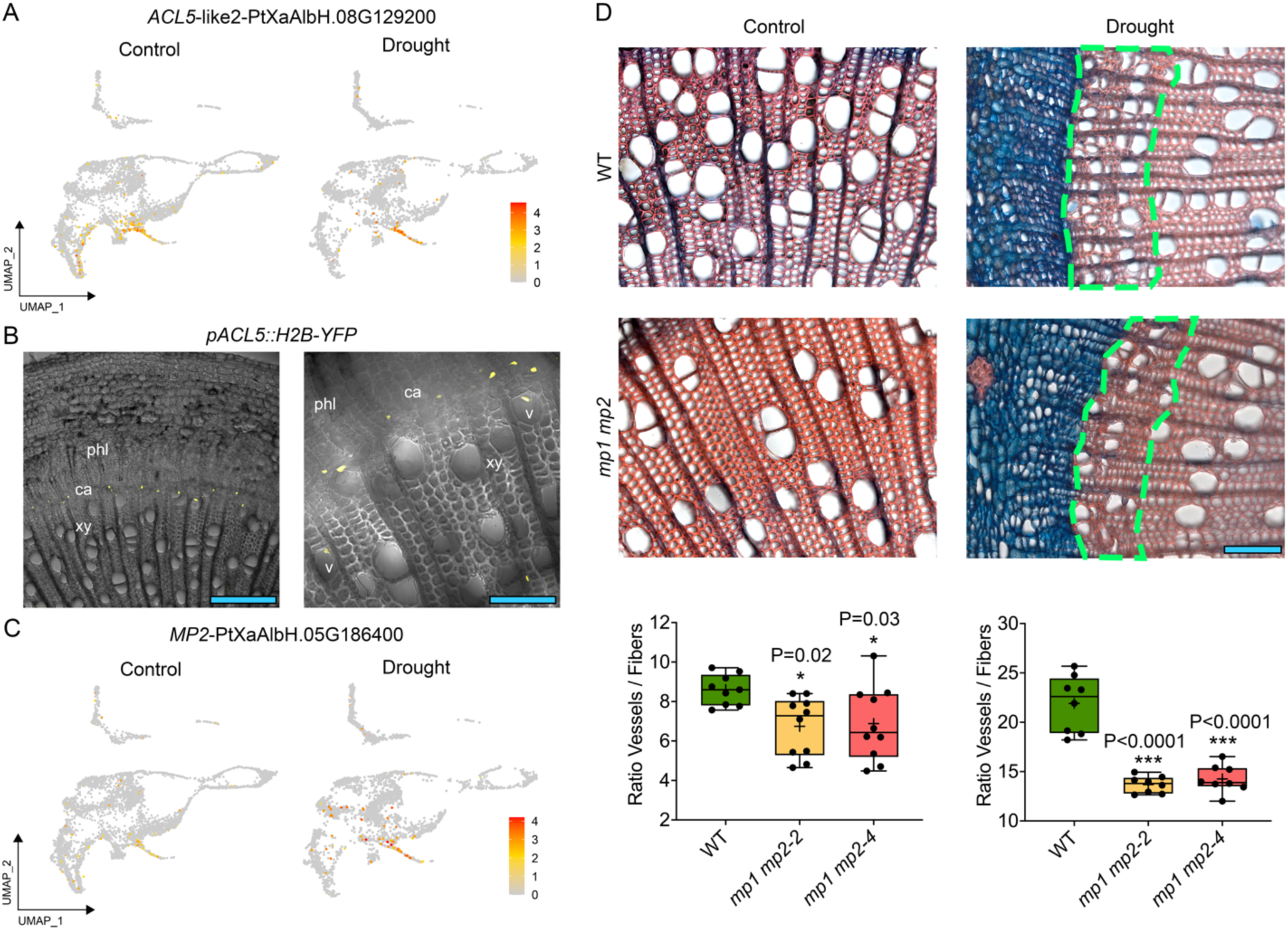
Functional characterization of hybrid poplar *MP* genes under well-watered and severe drought conditions. **A** *ACL5*-like 2 expression in the reclustering of cells related to xylem development under well-watered and drought conditions. **B** *H2B-YFP* fluorescent reporter activity under the *ACL5*-like 2 promoter indicated that it is mainly expressed at VC-PC and developing vessels. **C** *MP2* expression is induced and highly coexpressed with *ACL5*-like 2 under drought. **D** Under well-watered conditions, both *mp1 mp2* mutant lines showed a significant reduction in the vessel-to-fiber ratio in the secondary xylem compared to the wild type. This difference was more pronounced after 20 days under severe drought at the secondary xylem generated under drought stress (green dashed line). For each box-and-whisker plot, the center black line and the “+” represent the median and the mean, respectively; the box extends from the 25th to 75th percentiles; the whiskers are the maximum and the minimum values (n=8 to 10 trees). *One-way* ANOVA followed by Dunnett’s test determined statistical differences. Scale bars 200 μm and 80 μm in B at the left and right images, respectively, and 100 μm in D.

### Comparison of the xylem developmental trajectory under well-watered and severe drought conditions

Next, we used two alternative approaches to uncover divergent regulatory mechanisms in secondary xylem formation in well-watered and drought-treated plants. We inferred and compared the developmental trajectory of secondary xylem lineages in both conditions based on pseudotime analysis^38^. We also identified differentially expressed genes (DEGs) in each cell type involved in xylem formation between samples grown under well-watered and drought conditions.

To uncover regulatory mechanisms involved in xylem formation under well-watered and drought conditions, we inferred trajectories from clusters associated with vessel element differentiation: VC (cluster 5), PC (cluster 2), XMC/DX enriched in early differentiating vessels (cluster 1), SCW (cluster 3) and SCW/PCD (cluster 6) (Fig. 3A). We applied Slingshot^38^ to infer the trajectory based on ordering of cells along a pseudotime. As expected, the trajectory containing cells from both well-watered and drought samples originated in cluster VC, followed by PC and DX/XMC. At this point, the trajectory bifurcated to SCW (cluster 3) in one branch and SCW/PCD (cluster 6) in the other (Supplementary Fig. 6). The trajectory from cluster 5 to cluster 3 is referred to as Lineage 1, and from cluster 5 to cluster 6 as Lineage 2 hereafter (Supplementary Fig. 6). Next, we applied a model implemented in tradeSeq^39^ to identify genes with significant expression changes along these lineages, for each condition separately.

Under well-watered conditions, 5,024 DEGs were detected along Lineage 1, and 2,987 DEGs along Lineage 2 (FDR < 0.05) (Supplementary Table 2). Fewer DEGs were observed under drought: 1,043 genes were differentially expressed in Lineage 1, and 559 in Lineage 2 (Supplementary Table 2). Among the 1,043 genes differentially expressed under drought in Lineage 1, 1,023 (98%) were also identified as DEG in the well-watered condition (Supplementary Table 2). Thus, 4,001 genes were specific to the well-watered and only 20 genes to the drought conditions (Supplementary Table 2). A similar pattern was detected for Lineage 2, where 559 DEGs were observed under drought, of which 537 (96%) were shared with the well-watered condition. For Lineage 2, 2,450 genes were exclusive to the well-watered, and only 22 to drought (Supplementary Table 2).

To visualize gene expression patterns during xylem differentiation, we divided developmental trajectories into 30 equal bins, each containing an equal number of cells. Heatmaps of conserved DEGs revealed strikingly similar gene expression patterns across well-watered and drought conditions along both lineages (Supplementary Fig. 7). Interestingly, the 4,001 and 2,450 DEGs identified exclusively in well-watered Lineage 1 and Lineage 2, respectively, were also expressed under drought, displaying similar patterns despite lacking statistical significance (Supplementary Fig. 8). A similar trend was observed for the 20 and 22 DEGs unique to drought in both lineages (Supplementary Fig. 9).

In conclusion, comparing trajectories under well-watered and drought conditions revealed that, after 14 days of severe drought, all stages of xylem development still occur, though the process is slower under drought. This is supported by a lower proportion of PC under drought (12% in well-watered vs. 5% in drought conditions) and a higher representation of drought cells in early developmental stages (Supplementary Fig. 10). While genes involved in xylem differentiation were identified, the results suggest that wood plasticity is controlled by a similar gene regulatory network in both conditions, with altered expression patterns under drought. Specifically, genes detected only in the well-watered condition show similar expression trends under drought, though with slight variations causing non-significance. Often, these genes are activated later in the drought trajectory (Supplementary Fig. 8). This prompted a focus on DEGs within each cell type involved in secondary xylem formation to uncover the specific gene expression changes responsible for wood plasticity under drought.

### Cell-type specific reprogramming for wood plasticity under severe drought

To dissect the gene expression reprogramming that leads to wood plasticity under drought, we identified the DEGs across conditions in each cluster (Fig. 3A). Genes with |log2FC| > 0.35^40^ and FDR < 0.05 were considered DEGs. We identified the following numbers of DEGs: 383 in VC (121 up, 262 down); 1035 in PC (394 up, 641 down); 2907 in DX/XMC cluster 1 (1181 up, 1726 down); 3066 in DX/XMC cluster 0 (1277 up, 1789 down); 1065 XP (339 up, 726 down); 935 in SCW (332 up, 603 down); 1 down in SCW/PCD. (Fig. 3C and Supplementary Table 3).

We conducted Gene Ontology (GO) enrichment analyses to identify Biological Processes (BP) affected within the cell types involved in vessel formation; these are VC (cluster 5), Proliferating Cells (cluster 2), XMC/DC enriched in early differentiating vessels (cluster 1) and SCW (cluster 3). Among upregulated genes under drought, the BP response to water deprivation was enriched in the four clusters, pointing to a general response to the treatment. The response to abscisic acid and circadian rhythm were enriched in three clusters (PC, DX/XMC, SCW, and VC, DX/XMC, SCW). Between more cell type-specific responses, we found jasmonic acid signaling pathway, brassinosteroid homeostasis or auxin export, at the VC; negative regulation of DNA recombination and response to hydrogen peroxide at VC and PC; response to auxin, response to jasmonic acid or xylem development at XMC/DX; positive regulation of brassinosteroid signaling pathway and cell wall biogenesis at XMC/DX and SCW (Fig. 3D) (Supplementary Table 4).

In XMC/DX cluster 1, genes related to auxins included poplar homologs of *AUXIN RESPONSE FACTOR 5* (*ARF5*, *MP*), *ARF2*, *ARF8*, *AUXIN RESISTANT 1* (*AUX1*), *PIN-FORMED 1* (*PIN1*), and *AUXIN SIGNALING F-BOX 2* (*AFB2*). Among genes linked to xylem development, we identified *LONESOME HIGHWAY* (*LHW*), suggesting activation of the MP pathway under drought. Additionally, a significant number of circadian genes were induced, including homologs of *GIGANTEA* (*GI*) and *PSEUDO-RESPONSE REGULATORS* (*PRR2/5/7*), indicating that part of the gene expression reprogramming may be regulated by the circadian clock in these cells.

The list of downregulated genes showed general responses, such as translation, enriched in all the clusters. Between the cluster-specific enriched BP, we found auxin export across the plasma membrane at PC, regulation of cell growth at XMC/DX cluster 1, or autophagy at SCW (Fig. 3D) (Supplementary Table 5).

Our analysis revealed extensive gene expression reprogramming in secondary xylem formation, highlighting changes in auxin homeostasis and the circadian clock as key mechanisms driving anatomical adaptations under severe drought.

### The auxin signaling pathway regulates secondary xylem plasticity during severe drought

Polar auxin transport (PAT) regulates vessel expansion and patterning in the secondary xylem. To identify genes involved in wood plasticity under drought, we focused on two enriched biological processes related to auxins: response to auxin and auxin export across the plasma membrane. The response to auxin was enriched in upregulated genes in XMC/DX cluster 1, with *MP* (PtXaAlbH.05G186400) being the most induced *ARF* (Supplementary Table 3). Auxin export was enriched in downregulated genes in cluster 2 (PC), including two hybrid poplar copies of the *Arabidopsis* homolog *WALLS ARE THIN 1* (*WAT1*), designated as *WAT1* (PtXaAlbH.02G025500) and *WAT2* (PtXaAlbH.05G183400).

The expression of poplar *MP* under severe drought highly correlates with *ACL5*-like 2 (Fig. 4A-C). In Arabidopsis, *MP* is required for xylem identity in the root^41^. In poplar, both *MP* copies are mainly expressed in the procambium during primary vasculature development, and their expression significantly decreases as the vascular system matures, transitioning from primary to secondary xylem^42^. The secondary xylem develops more fibers as the plant matures to support the structure. We hypothesized that *MP* induction in the secondary xylem under drought may explain the higher vessel ratio observed. We generated CRISPR/Cas9 homozygous mutants for both *MP* copies (*mp1 mp2-2* and *mp1 mp2-4*), resulting in truncated protein versions (Supplementary Fig. 11). We grew nine plants per genotype for six weeks and examined internode 15 of the stem under well-watered and severe drought conditions. In the mutants, the vessel-to-fiber ratio in the secondary xylem was significantly reduced compared to the WT under well-watered conditions (22% and 20%, respectively). These differences were enhanced under severe drought (37% and 34%) (Fig. 4D). These findings suggest that *MP* induction during severe drought is crucial for maintaining proper vessel density in the secondary xylem.

The *WAT1* gene is required for fibers’ secondary cell wall formation in *Arabidopsis*^43,44^. Under well-watered conditions, hybrid poplar copies of *WAT1* are mainly expressed in the VC, PC, XP, and XMC cluster 1 (Fig. 5A). Under drought, *WAT1* is restricted to XMC cluster 1 and XP, while *WAT2* is expressed in these cells and at VC. Both copies were significantly downregulated at PC under severe drought (Supplementary Table 3). We hypothesize that poplar *WAT1* genes mediate auxin-driven regulation of secondary xylem development. We generated CRISPR/Cas9-mediated homozygous mutants (Supplementary Fig. 12). Growth measured as height and stem diameter was severely compromised at the double mutants. The *wat2* single mutant (Supplementary Fig. 13) height and stem diameter displayed values between WT and double mutant lines, suggesting *WAT* genes show additive functions in hybrid poplar development (Fig. 5B and Fig. 6B). At the secondary xylem, vessel element distribution was dramatically altered in the double mutant lines, showing entire radial lines of very narrow vessels (Fig. 5C). The vessel-to-fiber ratio was markedly elevated at both double mutant lines (*wat1 wat2-1 and wat1 wat2-9*) compared to the WT (181% and 151%, respectively) (Fig. 5C, D). Unlike *Arabidopsis*, fiber cell walls did not exhibit evident alterations (Fig. 5C).

**Fig. 5.**
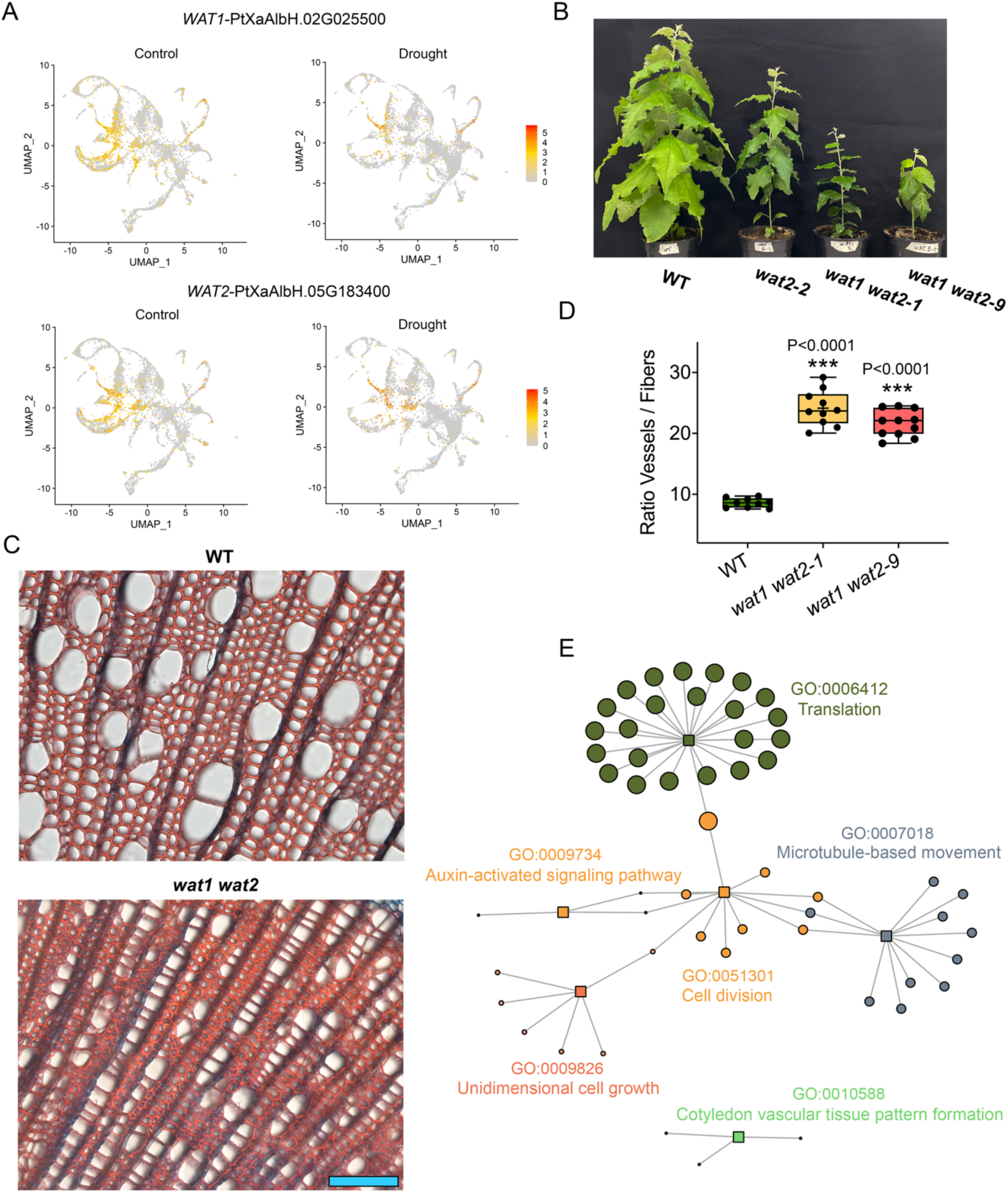
Characterization of hybrid poplar *WAT* genes function in secondary xylem. **A** *WAT1 and WAT2* expression in the overall dataset under well-watered and drought conditions. **B** After six weeks of growth in soil, growth was severely compromised in the *wat1 wat 2* double mutants, while single mutant phenotype remained between WT and double mutant lines. **C** Images of a stem cross-section of internode 15 of a WT and *wat1 wat2* mutant trees under well-watered conditions. At the secondary xylem, vessel element distribution was dramatically altered in the double mutant lines, with entire radial lines of very narrow vessels. **D** The vessel-to-fiber ratio was markedly elevated at both double mutant lines compared to the WT. For each box-and-whisker plot, the center black line and the “+” represent the median and the mean, respectively; the box extends from the 25th to 75th percentiles; the whiskers are the maximum and the minimum values (n=9 trees). *One-way* ANOVA followed by Dunnett’s test determined statistical differences. **E** GO network of enriched BPs in the list containing the top coexpressed genes with *WAT1* and *WAT2* in the overall dataset clustering. The network was built based on GO-gene and GO-GO relationships. The genes are represented as circles and GOs as squares. Circle size reflects the proportion of genes in each enriched GO. Scale bar 100 μm in D.

**Fig. 6.**
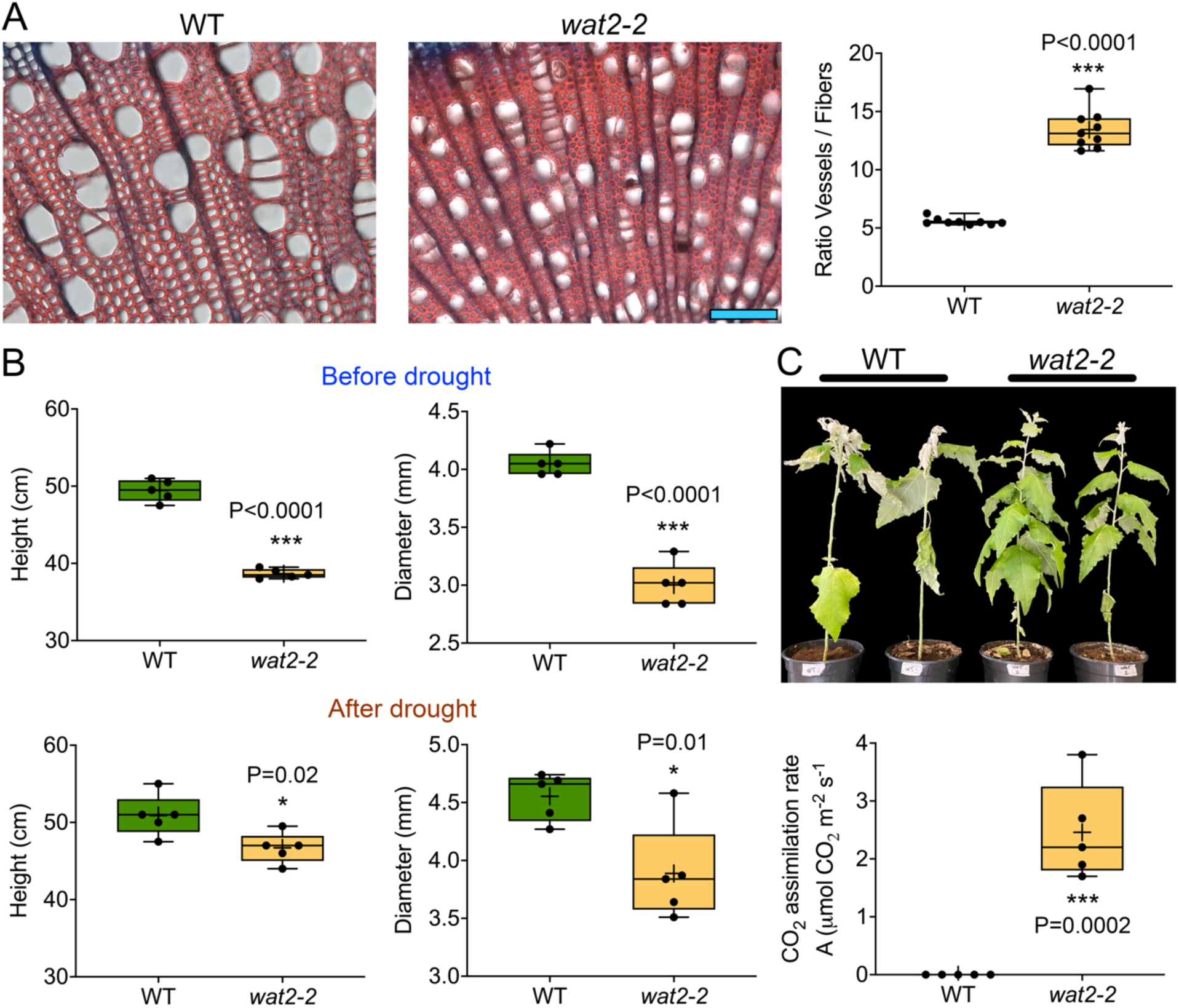
Characterization of the *wat2* mutant line adaptation to severe drought. **A** Images of a stem cross-section of internode 15 of a WT and *wat2* single mutant trees under well-watered conditions. At the secondary xylem, vessel element distribution was altered in the single mutant line, showing a higher vessel-to-fiber ratio and narrower vessels. For each box-and-whisker plot, the center black line and the “+” represent the median and the mean, respectively; the box extends from the 25th to 75th percentiles; the whiskers are the maximum and the minimum values (n=9 trees). Student’s t-test determined statistical differences. **B** Box plot showing height and diameter growth in WT and *wat2* mutant trees under well-watered conditions and after the severe drought treatment. For each box-and-whisker plot, the center black line and the “+” represent the median and the mean, respectively; the box extends from the 25th to 75th percentiles; the whiskers are the maximum and the minimum values (n=5 trees). Student’s t-test determined statistical differences. **C** After 20 days under severe drought, CO_2_ assimilation was significantly higher in *wat* trees than WT, indicating better fitness. Scale bar 100 μm in A.

We used hdWGCNA^45^ to generate the gene expression correlation network using whole stem clustering (Fig. 2A). We identified the 200 genes with the highest correlation scores with *WAT1* and *WAT2*. Of the 200 genes for each *WAT*, 135 were common for both (Supplementary Table 6). GO network analysis showed that these genes are involved in cell division, translation, unidimensional cell growth, auxin-activated signaling pathway or cotyledon vascular tissue pattern formation (Fig. 5E and Supplementary Table 7). Our results suggest that *WAT* genes in hybrid poplar are critical regulators of auxin homeostasis required for cell division and proper distribution of vessels in the secondary xylem by mediating the very early cell fate determination during (or early after) the vascular cambium cell division.

### The modification of secondary xylem anatomy drives physiological responses under severe drought

It is believed that vascular plants with smaller vessels can withstand lower water potential, thereby preventing xylem embolism under severe drought conditions^6^. However, there are very few examples of genetically modified trees with altered xylem development and better adaptation under drought. Transgenic poplars overexpressing *PtoERF15* exhibited a smaller vessel lumen area and more vessel cells than WT, resulting in drought tolerance^46^. Similarly to *PtoERF15*-OE lines^46^, our *wat2* mutant exhibited a 22% reduction in plant height and a 26% decrease in stem diameter compared with WT under well-watered conditions (Fig. 6A, B). In addition, the *wat2* mutant exhibited a significant increase of vessels-to-fibers ratio (142 %) and narrower vessels compared with WT (Fig. 6A). As the soil FC decreased to 15%, WT exhibited signs of impending death at the young stem, and leaves began to wilt. The leaves of the *wat2* line showed slight drooping. After 20 days under an FC of 15%, the physiological indicator associated with drought stress, the CO_2_ assimilation was significantly higher in the *wat2* trees compared with WT, indicating increased drought resilience (Fig. 6C). At the end of the treatment, the differences of plant height and diameter growth observed under well-watering conditions (22% and 26% respectively) were decreased to the levels of 8% and 15% respectively indicating a better severe drought tolerance of *wat2* mutants to the treatment (Fig. 6B). These results collectively suggest that the transcriptional regulation of *WAT* genes under severe drought play an active role in regulating drought tolerance and water transport in hybrid poplar.

## DISCUSSION

scRNA-seq has been used in several studies on the woody stems of various poplar species^34,42,47–50^. These analyses revealed significant differences in the proposed cell lineages for vessels, fibers, and xylem parenchyma. Four different models were suggested^34,47–49^, influenced by the differences in cell type annotation^51^. By snRNA-seq to provide the first cell type-specific characterization of gene expression reprogramming in poplar stems under severe drought conditions. Our data revealed two distinct groups of cells corresponding to the XMC/DX and another separate cluster for XP, 3A). The expression of previously identified marker genes for early differentiating vessels in *Arabidopsis*^37^ was enriched in one of these XMC/DX clusters (cluster 1, Fig. 3 A, B). Validation of hybrid poplar *ACL5*-like 2, expressed explicitly in a subdomain of the XMC/DX cluster 1 (Fig. 4A), indicated that this cluster contains cells undergoing early vessel differentiation. Our data show that from the XMC/DX clusters, the mature vessels and fibers formation converge in the cluster containing cells undergoing SCW formation, consistent with previous works^34,42^.In *Arabidopsis*, mature fibers and vessels are found in different clusters^37^. This finding reveals significant differences in SCW formation between tree mature stems and *Arabidopsis* roots that undergo secondary growth. Taken together, our data support the model proposed by Tung et al.^49^ regarding xylem development differentiation. Starting from the vascular cambium, the trajectory diverges into the ray parenchyma and the precursors of vessels and fibers.

We observed that hybrid poplar produced smaller yet more numerous vessels in the secondary xylem in response to severe drought. This finding aligns with previous studies involving different poplar species^13–15,17,52^. Our results suggest that auxin signaling is crucial in the trees’ wood plasticity. Under drought conditions, cambial activity is reduced, leading to severely suppressed radial growth^17^. Our findings indicate that after 14 days of severe drought, all stages of secondary xylem formation still occur, albeit at a diminished rate. In this regard, the arrest of cambial activity can be compared to a growth arrest during the onset of winter dormancy. It was initially believed that auxin levels in the cambium regulated growth arrest and dormancy in the VC. However, analyses in both active and dormant VC revealed very similar auxin levels in both states^53^. These findings suggest that other mechanisms, such as the cambium’s responsiveness to auxin or variations in active auxin distribution within the cells, underlie VC activity during growth-restricting conditions like dormancy and drought. A temporal transcriptomic analysis of poplar stems during dormancy highlighted two significant roles of auxins in the VC^54^: stimulating cambial cell division and preserving cambial meristem identity. Auxin levels in poplar stem remain unchanged during severe drought compared to the well-watered period^17^. In this study, we characterized the function of hybrid poplar *WAT1*-like genes. In *Arabidopsis, WAT1* facilitates auxin export from vacuoles to the cytoplasm^43,44^. The specific downregulation of both *WAT1* copies at PC and the limited growth observed in the *wat1 wat2* mutants highlights the critical role of *WAT1*-like genes in the auxin-mediated cell proliferation at the VC (Fig. 7).

**Fig. 7.**
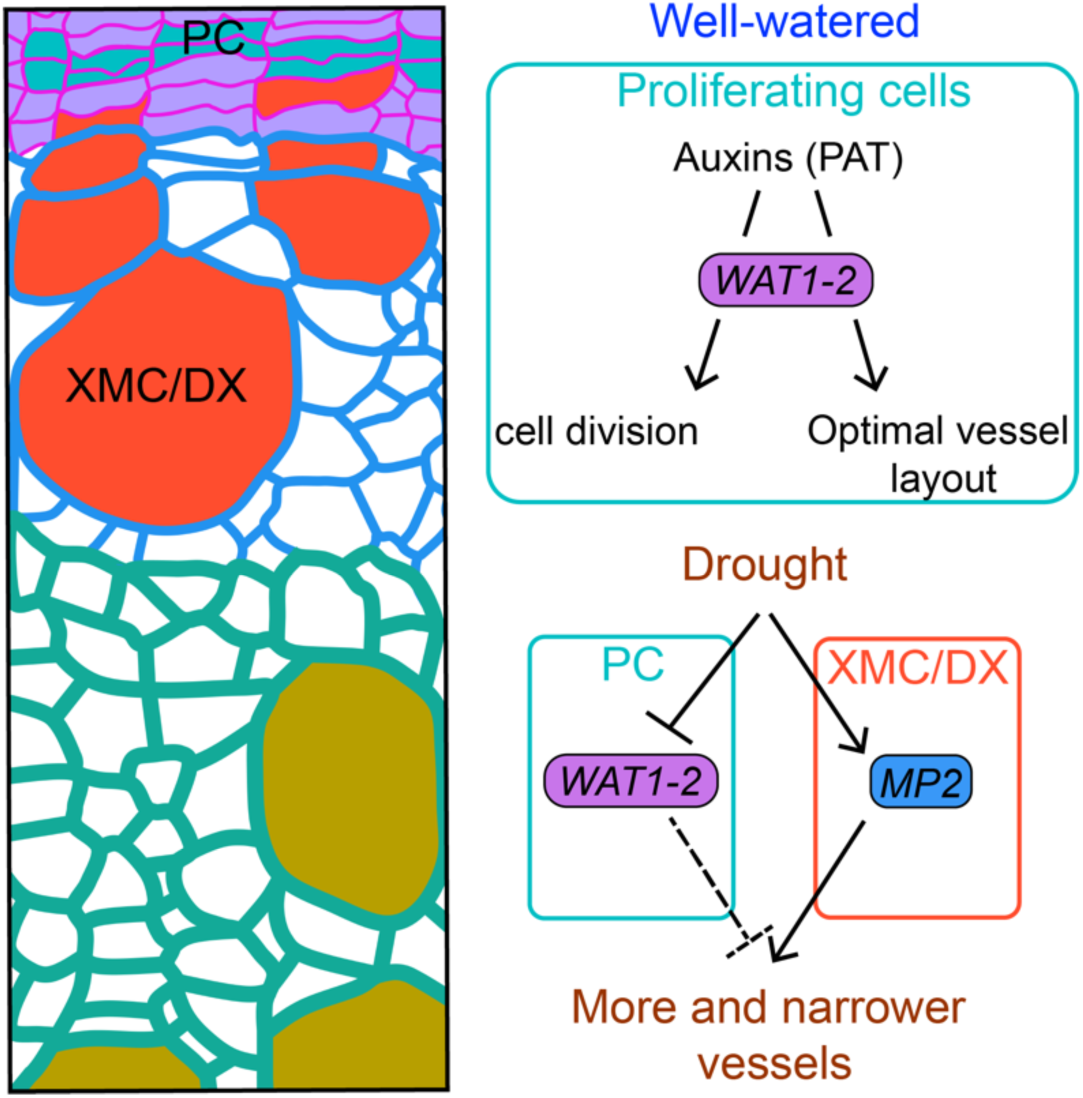
Graphical abstract depicting the functional characterization results of this study. Under well-watered conditions, hybrid poplar *WAT* genes regulate auxin-mediated cell division in the vascular cambium and coordinate the spatial patterning of vessels. During severe drought, the downregulation of *WAT* gene expression in the PC of the VC and the upregulation of the *MP2* gene in XMC/DX facilitate the development of secondary xylem with narrower and more abundant vessels.

Auxin establishes a gradient across the cambial zone, with the highest concentrations in the VC and their immediate xylem-side derivatives. Auxin levels diminish in the mature xylem and phloem^53,55,56^. PAT is essential for determining the proper spatial arrangement and size of vessels^57^. Inhibition of polar auxin transport results in the formation of vessels that are small in diameter, densely clustered, and angular in cross-section^57^. The *wat1 wat2* double mutant trees exhibit more and smaller vessels with a changed spatial pattern. These findings suggest that *WAT1*-like genes play a role in auxin-mediated cell fate determination in the xylem and imply that the downregulation of *WAT* genes expression under drought may lead to an increased number of vessels (Fig. 7).

Vessel dimensions and spatial distribution are critically important features of woody stems because they influence hydraulic efficiency and vulnerability to embolism, allowing the adaptation to various environments. However, our understanding of vessel developmental plasticity remains limited. In *Arabidopsis*, *WAT1* is required to form the SCW in fibers^43,44^. Our study reveals novel functions of hybrid poplar *WAT1*-like genes: regulating cell division in the vascular cambium and orchestrating the spatial patterning of vessels. Moreover, the *wat2* single mutant performed better under severe drought conditions than the WT, highlighting its potential for developing tree varieties with improved drought tolerance. Although the specific role of each *WAT1* gene copy in poplar will require future characterization, our observations of the *wat2* single mutant and *wat1 wat2* double mutant phenotypes suggest that they have additive functions in hybrid poplar development. Our findings also indicate that the induction and repression of auxin signaling genes contribute to the anatomical changes observed in secondary xylem under drought conditions. On one hand, the downregulation of *WAT1*-like genes results in more and narrower vessels. On the other hand, our results suggest that the upregulation of *MP* and likely other *ARFs* observed at the XMC/DX cluster 1 under severe drought may contribute to the formation of secondary xylem with an increased number of vessels under drought conditions (Fig. 7). These results imply a delicate balance of auxin cellular levels and signaling to create more functional vessels under severe drought. Our data also highlight the roles of various hormones regulating the drought response in the VC and secondary xylem under drought, such as gibberellins, jasmonic acid, abscisic acid, and brassinosteroids. Previous research indicates that jasmonic acid affects the architecture of secondary xylem under drought in poplar^46^.

Overall, our gene expression map of the hybrid poplar mature stem under well-watered and severe drought conditions at single-cell resolution provides a valuable resource for understanding how trees adapt to drought. In this study, we focused on analyzing the mechanisms of wood development at the single-cell level. Additionally, we compiled a table of all differentially expressed genes (DEGs) across the various cell types identified in the hybrid poplar stem in response to drought conditions, compared to the well-watered state (Fig. 2A) (Supplementary Table 8). This table will serve as a valuable resource for researchers investigating the cell type-specific responses of poplar stems to drought. We also conducted bulk RNA-seq, which confirmed the overall downregulation of hybrid poplar *WAT*1-like genes and the upregulation of *MP* and *GI* genes (Supplementary Table 9) and highlights the immense value of the spatial information provided by snRNA-seq data in identifying the molecular drivers of organ– and tissue-specific plasticity that enable plants to adapt to abiotic stresses.

## METHODS

### Plant material and growth conditions

Shoot tips and stem cuttings from in vitro-grown hybrid poplar (*Populus tremula*×*alba* INRA clone 717 1B4) were used as explants for plant multiplication. The explants were cut into small segments (10-15 mm long) and placed aseptically in Murashige and Skoog (MS) medium 1B (pH 5.7) supplemented with 2% sucrose and with indole acetic and indole butyric acids (0.5 mg/L) containing 0.7% (w/v) plant agar. Explants were grown under long-day (LD) 16-h light/8-h dark and 22°C conditions, 65% relative humidity, and 100 μmol m−2 s−1 photosynthetic photon flux. For the severe drought treatments, in vitro-cultivated hybrid poplars of WT, were transferred to pots containing peat, and grown under LD and 22°C conditions for 6 weeks.

Six weeks after potting, the plants were divided into two groups: well-watered (control) and severe drought treatment, with four biological replicates in each group. Irrigation was carefully controlled during 14 or 20 days of treatment. Soil moisture in the pot of each plant was measured daily with a TEROS 10 soil moisture probe attached to a ZL6 recording station (METER Group, USA). The treatments were performed similarly to those described previously^17^. Control plants were well-watered, keeping the solid moisture around FC (0.46 m^3^/m^3^) during the treatment. The weight of each pot was calculated at the FC before treatments. Severe drought treatment was achieved by stopping irrigation until the soil reached 15% of its field capacity for 14 days or 20 days. The wilting threshold was reached when soil moisture decreased to 0.10 m^3^/m^3^. Then, a small amount of water was added per pot daily to keep the 15% of FC by weight. Two independent experiments were performed to determine the anatomical changes of wood under severe drought.

### Wood Anatomical Analysis

Internode 15, where internode 1 refers to the stem portion between the first and the second fully extended leaves (still growing but not rolled up leaves), counting from the apex to the base, was fixed in 4% formaldehyde as described previously^58^. 40-60 µm thick cross-sections were obtained in a Leica VT1200S vibratome. Internodes were treated for 5 min in 50% ethanol and stained with 1.5% Safranine in 50% ethanol for 5 min. After three washes with distilled water (5 minutes each), sections were stained with the Alcian Blue Staining Solution (1 g of Alcian Blue, 97 ml of distilled water, and 3 ml of glacial acetic acid) for 5 min. After three washes with distilled water (5 minutes each), the sections were mounted on microscope slides with PBS. The images were obtained using a Zeiss LSM 880 microscope. The number of vessels and fibers of the eight trees subjected to the severe drought were measured in the portion of the secondary xylem generated before the treatment and in the secondary xylem developed under the severe drought (Fig. 1). At least two stem cross-sections per plant were measured using the ‘Analyze particles’ option of the ImageJ software.

### Nuclei isolation from hybrid poplar stem for single nucleus RNA-seq

To perform nuclei isolation for each library, internode 15 from one plant were placed on a glass plate with 400 μl of the Nuclei Extraction Buffer of the CyStain UV Precise P kit (Sysmex) supplemented with 2mM DTT, 0.8 U/μl of SUPERase-IN (Invitrogen), and 1:100 Protease Inhibitor (Sigma-Aldrich). Next, samples were chopped with a sterile razor blade for 2 min. This step was repeated with a 30-second interval. The homogenate was washed with 5 ml of the Staining Buffer of the CyStain UV Precise P kit from the glass plate and filtered through a pre-wetted (using the Staining Buffer) 40 μm strainer laid on top of a 50 ml conical tube placed on ice. To minimize the contamination of organelles such as chloroplast and remove the debris, the final solution, with around 5 ml of volume, was used to sort the nuclei using fluorescence-activated nuclei sorting. 80,000 nuclei were sorted with a total recovery volume of 100-120 μl in 1.5 ml RNase-free non-stick Eppendorf low binding tubes containing 10 μl of supplemented Nuclei Extraction Buffer as described above.

### Single nucleus cDNA and library preparation

45k nuclei were used to generate each snRNA-seq library following the GEXSCOPE Single Nucleus RNA library Kit protocol from Singleron Biotechnologies. Two snRNA-seq libraries for each treatment were generated.

### Quality control and cell clustering

The sequencing output was demultiplexed and processed with the software CeleScope v3.0.1 (Singleron), using the *Populus tremula*×*alba* 717-1B4 genome (v.5.1, HAP2) as reference. Then, using the raw counts matrix produced by CeleScope as input, the software EmptyDrops^59^, contained in the DropletUtils R package, was used to identify and remove instances where a barcode represented an empty well. The R package scDblFinder^60^ was used to identify doublets. The filtered dataset was loaded on Asc-Surat^18^, and only cells containing 400 and 200 or more expressed genes for the well-watered and drought libraries, respectively, were selected and used for the analysis.

### Cell clustering

Integration of the datasets from the replicates and cell clustering was performed using Asc-Seurat. The parameters used for cell clustering after integrating data from the four libraries (replicate 1 and replicate 2 for each treatment) are described in Supplementary Fig. 3A. The parameters used for the XDR data re-clustering are described in Supplementary Fig. 3B.

### Trajectory inference analysis

To compare the developmental trajectory of secondary xylem focusing on those cells enriched in vessel development, we used the Bioconductor packages Slingshot^38^, to infer the trajectories, and tradeSeq^39^, to identify genes whose expression changes along the trajectories. Using tradeSeq, a gene was considered differentially expressed within the trajectory if the Wald test, performed by the function *associationTest*, returned a False Discovery Rate smaller than 0.05.

### Weighted Correlation Network Analysis

Weighted Correlation Network Analysis (WGCNA) characterizes co-expression patterns in large gene expression datasets. We use the package hdWGCNA^45^, an adaptation of the original WGCNA^61^ method for single-cell datasets, to construct a co-expression network of genes expressed in the whole snRNA-seq dataset.

### Cluster-specific identification of differentially expressed genes between treatments

Differentially expressed genes under drought condition were identified using the FindMarkers function within the R package Seurat^62^, using the *bimod* test^63^ (|log2FC| ≥ 0.35 and p_val_adj < 0.05) (Table S3). The test was applied for each of the clusters involved in the secondary xylem development (Xylem Development Reclustering (XDR), Fig. 3A). Cluster-specific DEGs under drought were also identified for each cluster of the main clustered dataset (Fig. 2A) by the same method (Supplementary Table 8).

### Gene Ontology Enrichment Analysis

Gene Ontology enrichment analysis to identify enriched biological processes was performed with topGO in R, using the algorithm *weight01*, using the list of DEGs in clusters 1, 2, 3, and 5 of the Xylem Development Reclustering (XDR, Fig. 3A).

### Validation of hybrid poplar *ACL5* expression by promoter-fluorescent reporter fusion

The MultiSite Gateway Three-Fragment Vector Construction kit was used to clone the DNA constructs. A 2057 bp segment of the promoter (up to the ATG start codon) of the hybrid poplar (HAP1) *ACL5* (PtXaAlbH.08G129200) gene was amplified by PCR. The promoter sequence was cloned by the BP reaction into the pDONR P4-P1R vector. The same strategy was applied to clone the fluorescent reporter H2B-YFP into the pDONR 221. Finally, the LR reaction assembled the transcriptional unit into the destination vector using an empty version of the pDONR P2R-P3 for position 3 of the system. We generated stable transgenic lines. *Agrobacterium tumefaciens*-mediated transformation was performed using the strain GV3101 in the *Populus tremula*×*alba* 717-1B4 genotype, using a previously developed protocol^64^. We used the antibiotic kanamycin to identify the positive events of the transformation.

Three-week-old *in vitro* transgenic plants containing the *pACL5::H2B-YFP* construct were transferred to pots. After 4 weeks of growth under a long-day regimen at 22°C, 65% relative humidity and 100 μmol m−2 s−1 photosynthetic photon flux, a 5 mm-long portion of the apex, internode 15 was fixed in 4% formaldehyde as described previously^58^. 200 µm thick cross-sections were obtained in a Leica VT1200S vibratome. The sections were mounted on microscope slides with PBS and 50% glycerol. Images were obtained under the confocal microscope (Leica TCS SP8) at an excitation wavelength of 514 nm.

### Functional characterization of *MP*– and *WAT1*-like genes in hybrid poplar

#### Cloning

Hybrid poplar CRISPR/Cas9-mediated mutants for *MP* and *WAT1* genes were generated expressing three and two single guide RNAs (sgRNAs), respectively, targeting the two hybrid poplar paralog genes coding sequence (Supplementary Fig. 11, Supplementary Fig. 12 and Supplementary Fig. 13). The DNA constructs were generated using the Golden Gate-based system that we designed for multi-site genome editing in hybrid poplar^65^. *Agrobacterium*-mediated transformation was performed using the strain GV3101 in the *Populus tremula*×*alba* 717-1B4 genotype, as described above. Independent lines obtained by kanamycin selection were genetically screened to identify the specific allele mutation by Nanopore sequencing of a PCR fragment spanning CRISPR/Cas9 target sites of the genes of interest (Plasmidsaurus company) (Supplementary Fig. 11, Supplementary Fig. 12 and Supplementary Fig. 13).

#### Characterization of the secondary xylem mutant lines

Wood anatomy was analyzed as described above. After 6 weeks of growth under a long-day regimen at 22°C, 65% relative humidity, and 100 μmol m−2 s−1 photosynthetic photon flux, the WT and *mp1 mp2* mutant lines were subjected to control or severe drought for 20 days as described above. The secondary xylem anatomy of *wat1 wat2* and *wat2* mutant trees were evaluated under well-watered conditions. The vessel-to-fiber ratio was determined as described above.

### RNA-seq

Total RNA extraction was performed as described previously^66^ from the stem of three well-watered and three severe drought-treated trees (three biological replicates per treatment). RNA was sent to Macrogen company (South Korea). The RNA-seq libraries were generated using the TruSeq Stranded mRNA Library Prep Kit and sequenced on the Illumina NovaSeq X platform. Reads were then mapped to the reference genome of *Populus tremula*×*alba* INRA clone 717 1B4, HAP2 v5.1 with HISAT2. DEGs were identified using the EdgeR package in R with the significance threshold set at FDR < 0.05.

## DATA AVAILABILITY

SnRNA-seq sequencing raw data, CeleScope outputs after filtering, and bulk RNA-seq raw data have been deposited in the NCBI Gene Expression Omnibus and are accessible through accession number GSE283835.

## ACKNOWLEDGMENTS

We thank Drs. Crisanto Gutiérrez Armenta and Bénédicte Desvoyes for their technical support with nuclei isolation and library preparation.

## CONTRIBUTIONS

Conceptualization: D.C.; Methodology: D.C., D.G.S., W.J.P.; Data analysis: D.C., W.J.P., D.G.S.; Validation: D.C., A.P.D.; Resources: D.C., I.A., M.P., C.D., M.K.; Data curation: W.J.P., D.G.S., I.A., M.P.; Writing – original draft: D.C., M.K., M.P.; Writing – review & editing: D.G.S., W.J.P., C.D., I.A., M.P., M.K.; Funding acquisition: D.C., M.K., I.A.

## REFERENCES

1. Senf, C., Buras, A., Zang, C. S., Rammig, A. & Seidl, R. Excess forest mortality is consistently linked to drought across Europe. Nat Commun 11, 6200 (2020).

2. Anderegg, W. R. L., Kane, J. M. & Anderegg, L. D. L. Consequences of widespread tree mortality triggered by drought and temperature stress. Nature Clim Change 3, 30–36 (2013).

3. Lorenzo, M. N. & Alvarez, I. Climate change patterns in precipitation over Spain using CORDEX projections for 2021–2050. Science of The Total Environment 723, 138024 (2020).

4. Ramirez, F. & Kallarackal, J. *Responses of Fruit Trees to Global Climate Change*. (Springer International Publishing, Cham, 2015). doi:10.1007/978-3-319-14200-5.

5. Choat, B. et al. Triggers of tree mortality under drought. Nature 558, 531–539 (2018).

6. Rodriguez-Zaccaro, F. D. & Groover, A. Wood and water: How trees modify wood development to cope with drought. Plants People Planet 1, 346–355 (2019).

7. Furuta, K. M., Hellmann, E. & Helariutta, Y. Molecular Control of Cell Specification and Cell Differentiation During Procambial Development. Annu. Rev. Plant Biol. 65, 607–638 (2014).

8. Spicer, R. & Groover, A. Evolution of development of vascular cambia and secondary growth. New Phytologist 186, 577–592 (2010).

9. Samuels, A. L., Kaneda, M. & Rensing, K. H. The cell biology of wood formation: from cambial divisions to mature secondary xylemThis review is one of a selection of papers published in the Special Issue on Plant Cell Biology. Can. J. Bot. 84, 631–639 (2006).

10. Agustí, J. & Blázquez, M. A. Plant vascular development: mechanisms and environmental regulation. Cell. Mol. Life Sci. 77, 3711–3728 (2020).

11. Ribeiro, C. L. et al. The uncharacterized gene *EVE* contributes to vessel element dimensions in *Populus*. Proc Natl Acad Sci USA 117, 5059–5066 (2020).

12. Fonti, P., Heller, O., Cherubini, P., Rigling, A. & Arend, M. Wood anatomical responses of oak saplings exposed to air warming and soil drought. Plant Biology 15, 210–219 (2013).

13. Arend, M. & Fromm, J. Seasonal change in the drought response of wood cell development in poplar. Tree Physiology 27, 985–992 (2007).

14. Beniwal, R. S., Langenfeld-Heyser, R. & Polle, A. Ectomycorrhiza and hydrogel protect hybrid poplar from water deficit and unravel plastic responses of xylem anatomy. Environmental and Experimental Botany 69, 189–197 (2010).

15. Fichot, R. et al. Xylem anatomy correlates with gas exchange, water-use efficiency and growth performance under contrasting water regimes: evidence from Populus deltoides x Populus nigra hybrids. Tree Physiology 29, 1537–1549 (2009).

16. Song, J. et al. The influence of nitrogen availability on anatomical and physiological responses of Populus alba × P. glandulosa to drought stress. BMC Plant Biol 19, 63 (2019).

17. Yu, D. et al. Wood Formation under Severe Drought Invokes Adjustment of the Hormonal and Transcriptional Landscape in Poplar. IJMS 22, 9899 (2021).

18. Pereira, W. J. et al. Asc-Seurat: analytical single-cell Seurat-based web application. BMC Bioinformatics 22, 556 (2021).

19. Randall, R. S., et al. *AINTEGUMENTA* and the D-type cyclin *CYCD3;1* regulate root secondary growth and respond to cytokinins. Biology Open 4, 1229–1236 (2015).

20. Eswaran, G. et al. Identification of cambium stem cell factors and their positioning mechanism. Science 386, 646–653 (2024).

21. Kucukoglu, M., Nilsson, J., Zheng, B., Chaabouni, S. & Nilsson, O. *WUSCHEL – RELATED HOMEOBOX 4 ( WOX 4)* –like genes regulate cambial cell division activity and secondary growth in *Populus* trees. New Phytologist 215, 642–657 (2017).

22. Fisher, K. & Turner, S. PXY, a Receptor-like Kinase Essential for Maintaining Polarity during Plant Vascular-Tissue Development. Current Biology 17, 1061–1066 (2007).

23. Etchells, J. P. & Turner, S. R. The PXY-CLE41 receptor ligand pair defines a multifunctional pathway that controls the rate and orientation of vascular cell division. Development 137, 767– 774 (2010).

24. Muñiz, L. et al. ACAULIS5 controls *Arabidopsis* xylem specification through the prevention of premature cell death. Development 135, 2573–2582 (2008).

25. Ditengou, F. A. et al. Characterization of auxin transporter PIN 6 plasma membrane targeting reveals a function for PIN 6 in plant bolting. New Phytologist 217, 1610–1624 (2018).

26. Zhou, J., Zhong, R. & Ye, Z.-H. Arabidopsis NAC Domain Proteins, VND1 to VND5, Are Transcriptional Regulators of Secondary Wall Biosynthesis in Vessels. PLoS ONE 9, e105726 (2014).

27. Abbas, M. et al. Involvement of CesA4, CesA7-A/B and CesA8-A/B in secondary wall formation in Populus trichocarpa wood. Tree Physiology 40, 73–89 (2020).

28. Taylor-Teeples, M. et al. An Arabidopsis gene regulatory network for secondary cell wall synthesis. Nature 517, 571–575 (2015).

29. Avci, U., Earl Petzold, H., Ismail, I. O., Beers, E. P. & Haigler, C. H. Cysteine proteases XCP1 and XCP2 aid micro-autolysis within the intact central vacuole during xylogenesis in Arabidopsis roots. The Plant Journal 56, 303–315 (2008).

30. Wallner, E., Tonn, N., Shi, D., Jouannet, V. & Greb, T. *SUPPRESSOR OF MAX2 1-LIKE 5* promotes secondary phloem formation during radial stem growth. Plant J 102, 903–915 (2020).

31. Nguyen, V. P. et al. Identification and functional analysis of a promoter sequence for phloem tissue specific gene expression from Populus trichocarpa. J. Plant Biol. 60, 129–136 (2017).

32. Dinant, S. et al. Diversity of the Superfamily of Phloem Lectins (Phloem Protein 2) in Angiosperms. Plant Physiology 131, 114–128 (2003).

32. Kim, J.-Y. et al. Distinct identities of leaf phloem cells revealed by single cell transcriptomics. The Plant Cell 33, 511–530 (2021).

34. Chen, Y. et al. Transcriptional landscape of highly lignified poplar stems at single-cell resolution. Genome Biol 22, 319 (2021).

35. Rains, M. K., Gardiyehewa De Silva, N. D. & Molina, I. Reconstructing the suberin pathway in poplar by chemical and transcriptomic analysis of bark tissues. Tree Physiology 38, 340– 361 (2018).

36. Miguel, A., Milhinhos, A., Novák, O., Jones, B. & Miguel, C. M. The *SHORT-ROOT* –like gene *PtSHR2B* is involved in *Populus* phellogen activity. EXBOTJ 67, 1545–1555 (2016).

37. Lyu, M. et al. The dynamic and diverse nature of parenchyma cells in the Arabidopsis root during secondary growth. Preprint at 10.1101/2024.07.18.604073 (2024).

38. Street, K. et al. Slingshot: cell lineage and pseudotime inference for single-cell transcriptomics. BMC Genomics 19, 477 (2018).

39. Van den Berge, K., et al. Trajectory-based differential expression analysis for single-cell sequencing data. Nat Commun 11, 1201 (2020).

40. Chen, X. et al. Single-cell transcriptomic analysis of pea shoot development and cell-type-specific responses to boron deficiency. The Plant Journal 117, 302–322 (2024).

41. Smetana, O. et al. High levels of auxin signalling define the stem-cell organizer of the vascular cambium. Nature 565, 485–489 (2019).

42. Du, J. et al. High-resolution anatomical and spatial transcriptome analyses reveal two types of meristematic cell pools within the secondary vascular tissue of poplar stem. Molecular Plant 16, 809–828 (2023).

43. Ranocha, P. et al. Arabidopsis WAT1 is a vacuolar auxin transport facilitator required for auxin homoeostasis. Nat Commun 4, 2625 (2013).

44. Ranocha, P. et al. Walls are thin 1 (WAT1), an Arabidopsis homolog of Medicago truncatula NODULIN21, is a tonoplast-localized protein required for secondary wall formation in fibers: Tonoplastic WAT1 and secondary wall formation. The Plant Journal 63, 469–483 (2010).

45. Morabito, S., Reese, F., Rahimzadeh, N., Miyoshi, E. & Swarup, V. hdWGCNA identifies co-expression networks in high-dimensional transcriptomics data. Cell Reports Methods 3, 100498 (2023).

46. Kong, L. et al. The AP2 / ERF transcription factor PTOERF15 confers drought tolerance via JA –mediated signaling in *Populus*. New Phytologist 240, 1848–1867 (2023).

47. Li, H. et al. Single-cell RNA sequencing reveals a high-resolution cell atlas of xylem in *Populus*. J Integr Plant Biol 63, 1906–1921 (2021).

48. Xie, J., Li, M., Zeng, J., Li, X. & Zhang, D. Single-cell RNA sequencing profiles of stem-differentiating xylem in poplar. Plant Biotechnology Journal 20, 417–419 (2022).

49. Tung, C.-C. et al. Single-cell transcriptomics unveils xylem cell development and evolution. Genome Biol 24, 3 (2023).

50. Li, R., Wang, Z., Wang, J.-W. & Li, L. Combining single-cell RNA sequencing with spatial transcriptome analysis reveals dynamic molecular maps of cambium differentiation in the primary and secondary growth of trees. Plant Communications 4, 100665 (2023).

51. Chen, Y.-L. et al. Merit of integrating in situ transcriptomics and anatomical information for cell annotation and lineage construction in single-cell analyses of Populus. Genome Biol 25, 85 (2024).

52. Bogeat-Triboulot, M.-B. et al. Gradual Soil Water Depletion Results in Reversible Changes of Gene Expression, Protein Profiles, Ecophysiology, and Growth Performance in *Populus euphratica*, a Poplar Growing in Arid Regions. Plant Physiology 143, 876–892 (2007).

53. Uggla, C., Mellerowicz, E. J. & Sundberg, B. Indole-3-Acetic Acid Controls Cambial Growth in Scots Pine by Positional Signaling1. Plant Physiology 117, 113–121 (1998).

54. Baba, K. et al. Activity–dormancy transition in the cambial meristem involves stage-specific modulation of auxin response in hybrid aspen. Proc. Natl. Acad. Sci. U.S.A. 108, 3418–3423 (2011).

55. Uggla, C., Moritz, T., Sandberg, G. & Sundberg, B. Auxin as a positional signal in pattern formation in plants. Proc. Natl. Acad. Sci. U.S.A. 93, 9282–9286 (1996).

56. Tuominen, H., Puech, L., Fink, S. & Sundberg, B. A Radial Concentration Gradient of Indole-3-Acetic Acid Is Related to Secondary Xylem Development in Hybrid Aspen. Plant Physiology 115, 577–585 (1997).

57. Johnson, D. et al. Polar auxin transport is implicated in vessel differentiation and spatial patterning during secondary growth in *Populus*. Am J Bot 105, 186–196 (2018).

58. Conde, D., González-Melendi, P. & Allona, I. Poplar stems show opposite epigenetic patterns during winter dormancy and vegetative growth. Trees 27, 311–320 (2013).

59. participants in the 1st Human Cell Atlas Jamboree et al. EmptyDrops: distinguishing cells from empty droplets in droplet-based single-cell RNA sequencing data. Genome Biol 20, 63 (2019).

60. Germain, P.-L., Lun, A., Garcia Meixide, C., Macnair, W. & Robinson, M. D. Doublet identification in single-cell sequencing data using scDblFinder. F1000Res 10, 979 (2022).

61. Langfelder, P. & Horvath, S. WGCNA: an R package for weighted correlation network analysis. BMC Bioinformatics 9, 559 (2008).

62. Hao, Y. et al. Integrated analysis of multimodal single-cell data. Cell 184, 3573–3587.e29 (2021).

63. McDavid, A. et al. Data exploration, quality control and testing in single-cell qPCR-based gene expression experiments. Bioinformatics 29, 461–467 (2013).

64. Gallardo, F. et al. Expression of a conifer glutamine synthetase gene in transgenic poplar. Planta 210, 19–26 (1999).

65. Triozzi, P. M., Schmidt, H. W., Dervinis, C., Kirst, M. & Conde, D. Simple, efficient and open-source CRISPR/Cas9 strategy for multi-site genome editing in Populus tremula x alba. Tree Physiology.

66. Ibañez, C. et al. Overall Alteration of Circadian Clock Gene Expression in the Chestnut Cold Response. PLoS ONE 3, e3567 (2008).

