## Supplementary Figures for "Single-nuclei transcriptomics revealed auxin-driven mechanisms of wood plasticity and severe drought tolerance in poplar"

| | *Ratio Vessels/Fibers at the secondary xylem generated under control* | *Ratio Vessels/Fibers at the secondary xylem generated under drought* |
|---|---|---|
| *TREE 1* | 5.802 % | 15.852 % |
| *TREE 2* | 5.350 % | 15.575 % |
| *TREE 3* | 4.987 % | 12.994 % |
| *TREE 4* | 5.217 % | 16.897 % |
| *TREE 5* | 6.073 % | 15.411 % |
| *TREE 6* | 7.251 % | 17.815 % |
| *TREE 7* | 7.755 % | 17.538 % |
| *TREE 8* | 5.211 % | 21.701 % |

**Supplementary Fig. 1.** Measurements of the vessel ratio (defined as the proportion of vessels to fibers) at the secondary xylem formed during the well-watering period (control) and under severe drought stress (20 days under drought) in eight trees at the internode 15.

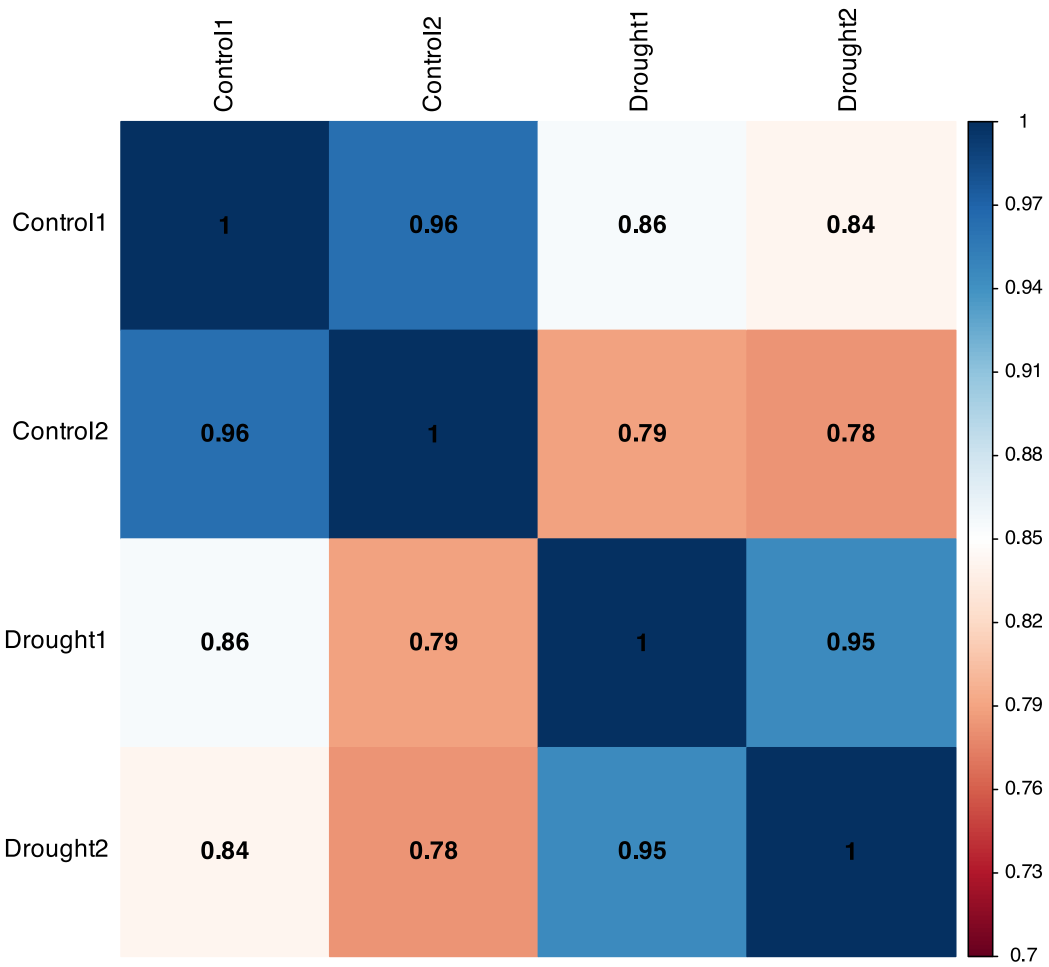


**Supplementary Fig. 2.** Pearson correlation of the average gene expression between the four snRNA-seq libraries generated, two for the control and two for the drought treatments.

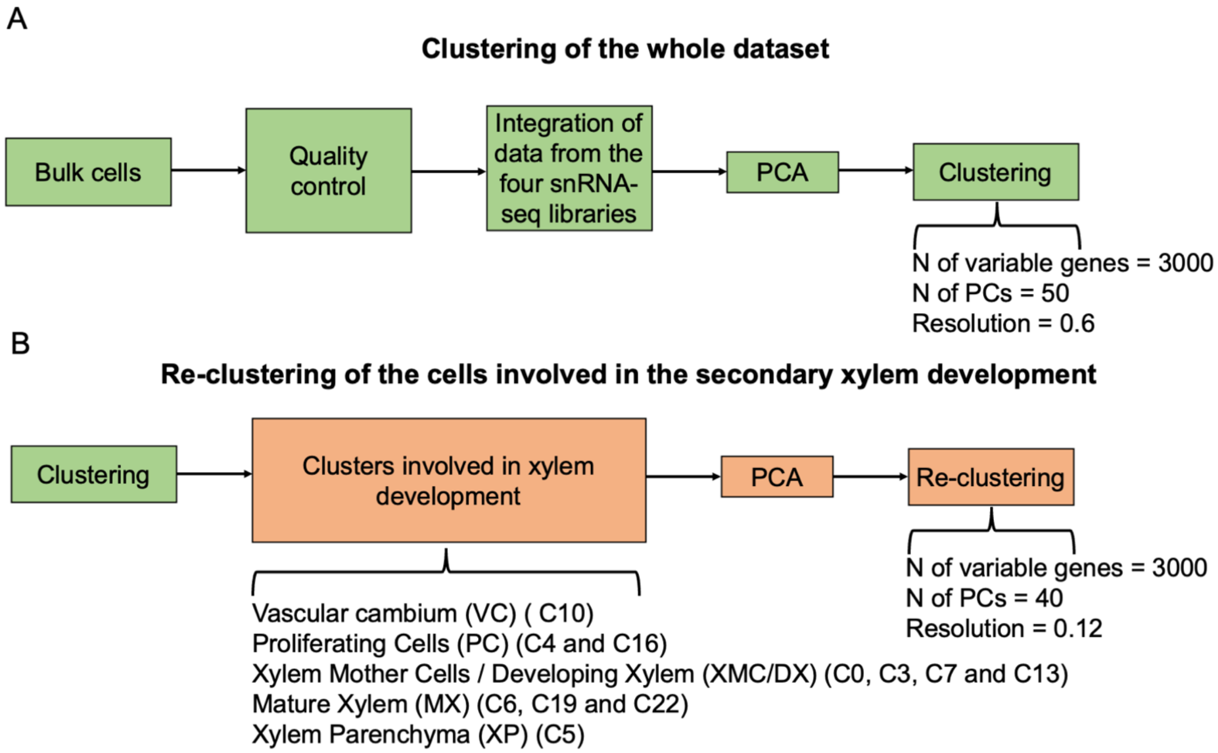


**Supplementary Fig. 3. A** Workflow followed to integrate and cluster the individual nuclei transcriptomes obtained from the hybrid poplar stem under well-watering and drought conditions. **B** Cells involved in the secondary xylem development were re-clustered following the parameters shown in this figure.

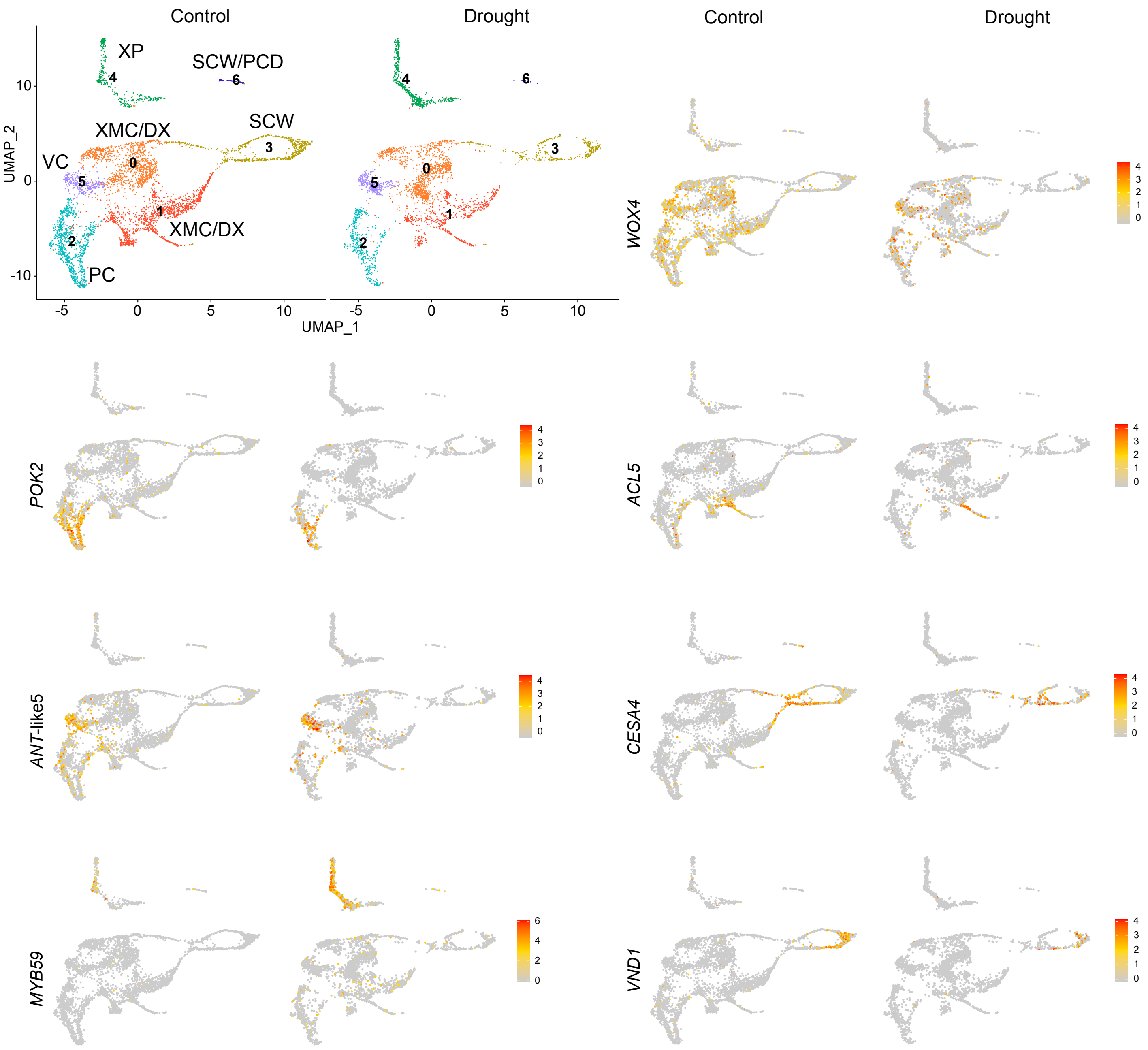


**Supplementary Fig. 4.** Visualization by UMAP of the hybrid poplar cell population involved in the secondary xylem development after the re-clustering. Dots, individual cells; color, cell clusters. Expression profiles of well-known markers identified proliferating cells (PC) (*POK2*-PtXaAlbH.04G113400), vascular cambium (VC) (*ANT*-like 5-PtXaAlbH.06G140900), xylem mother cells/developing xylem (XMC/DX) (*ACL5*-PtXaAlbH.08G129200), xylem parenchyma (XP) (*MYB59*, PtXaAlbH.01G196500), secondary cell wall (SCW) (*CESA4*-PtXaAlbH.11G050000, *VND1*-PtXaAlbH.07G013700). *WOX4*-PtXaAlbH.14G016600 is expressed at PC, VC, and XMC/DX.

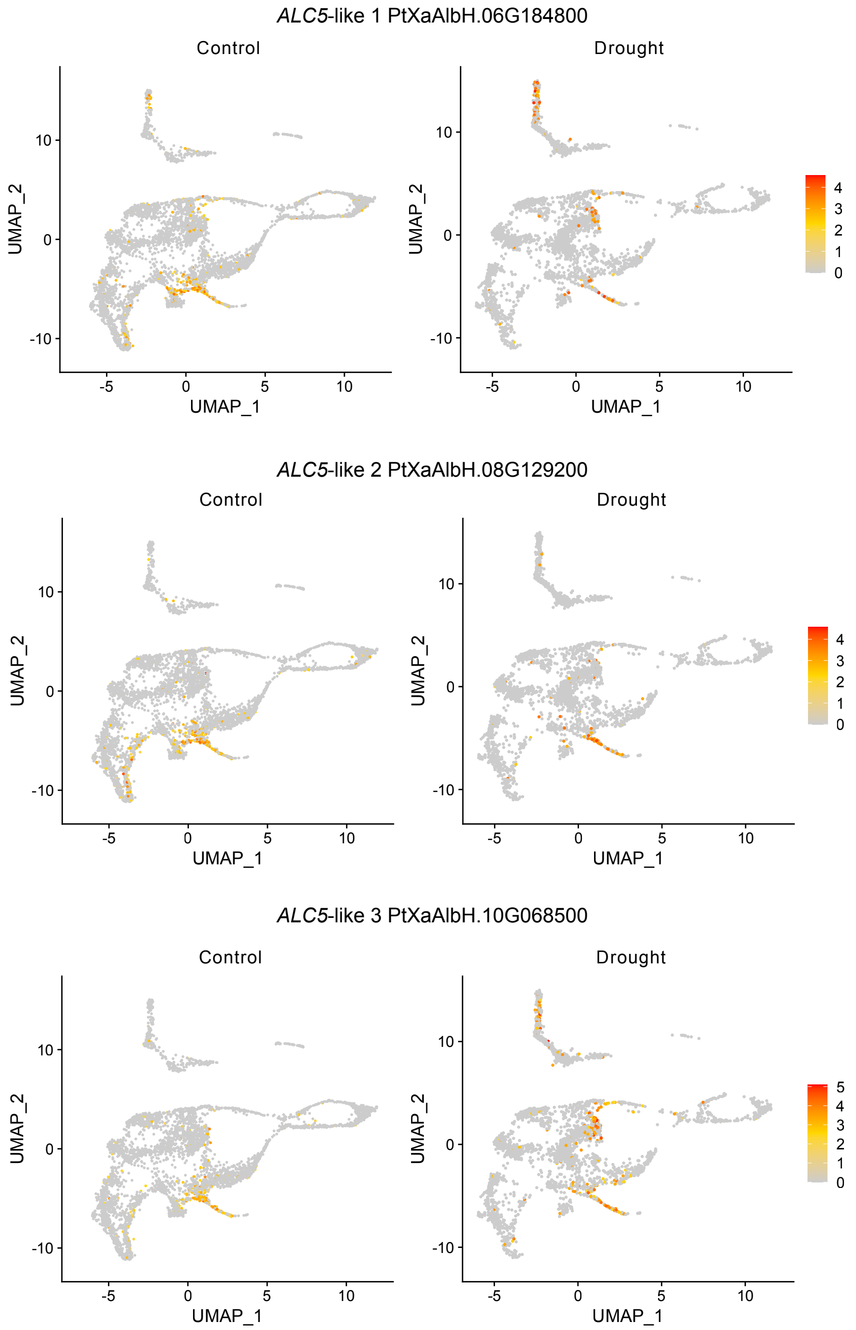


**Supplementary Fig. 5.** Expression of hybrid poplar *ACL5*-like genes in the cell population involved in the secondary xylem development.

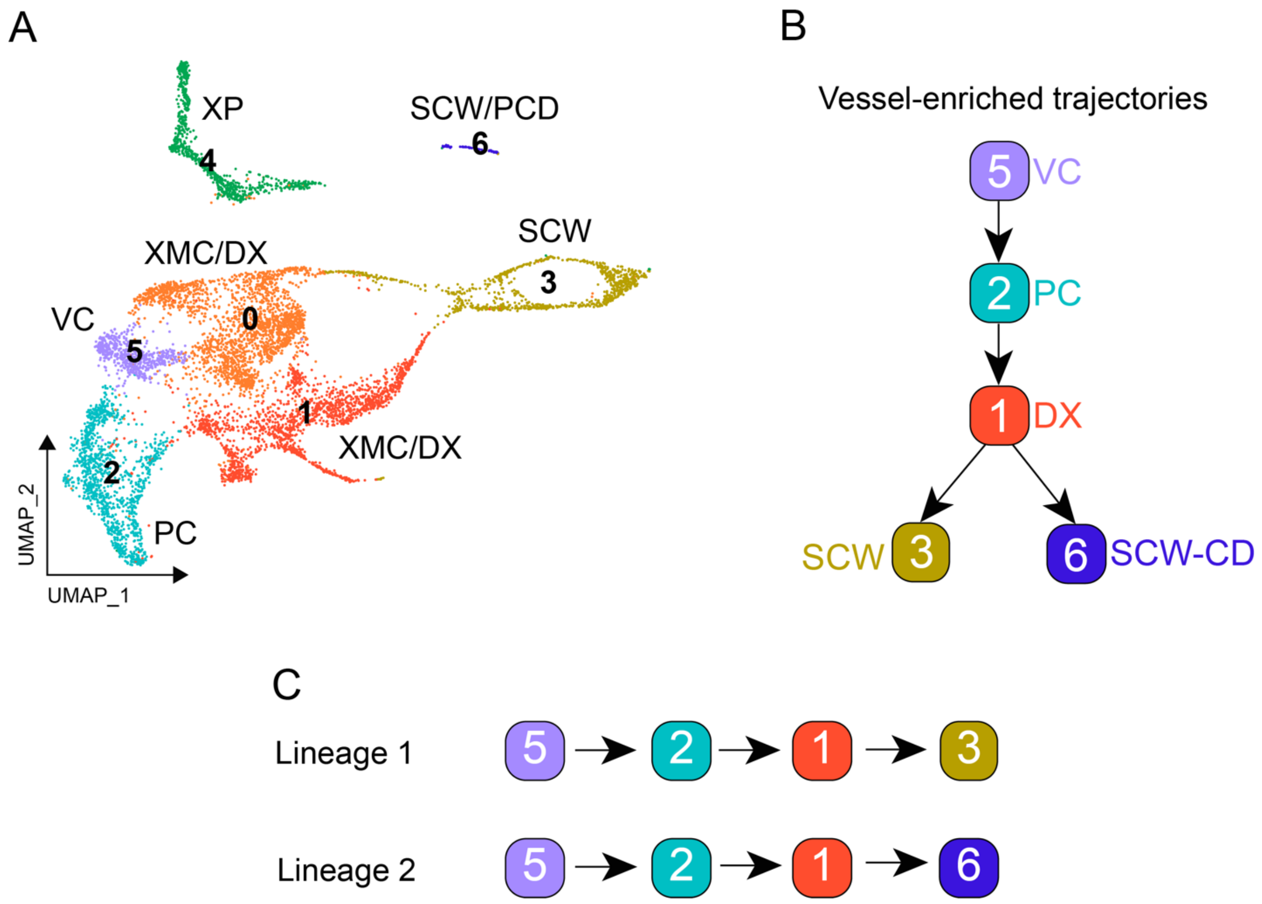


**Supplementary Fig. 6. A** Well-watered/drought integrated data visualization by UMAP of the hybrid poplar cell population involved in the secondary xylem development after the re-clustering. Dots, individual cells; color, cell clusters. **B** Slingshot was used in this re-clustered dataset to generate the trajectory for vessel development. **C** The trajectory bifurcates from cluster 1, originating two lineages that we named Lineage 1 and Lineage 2. This trajectory was also used in tradeSeq to identify genes whose expression significantly changes along the trajectories under well-watered and drought conditions.

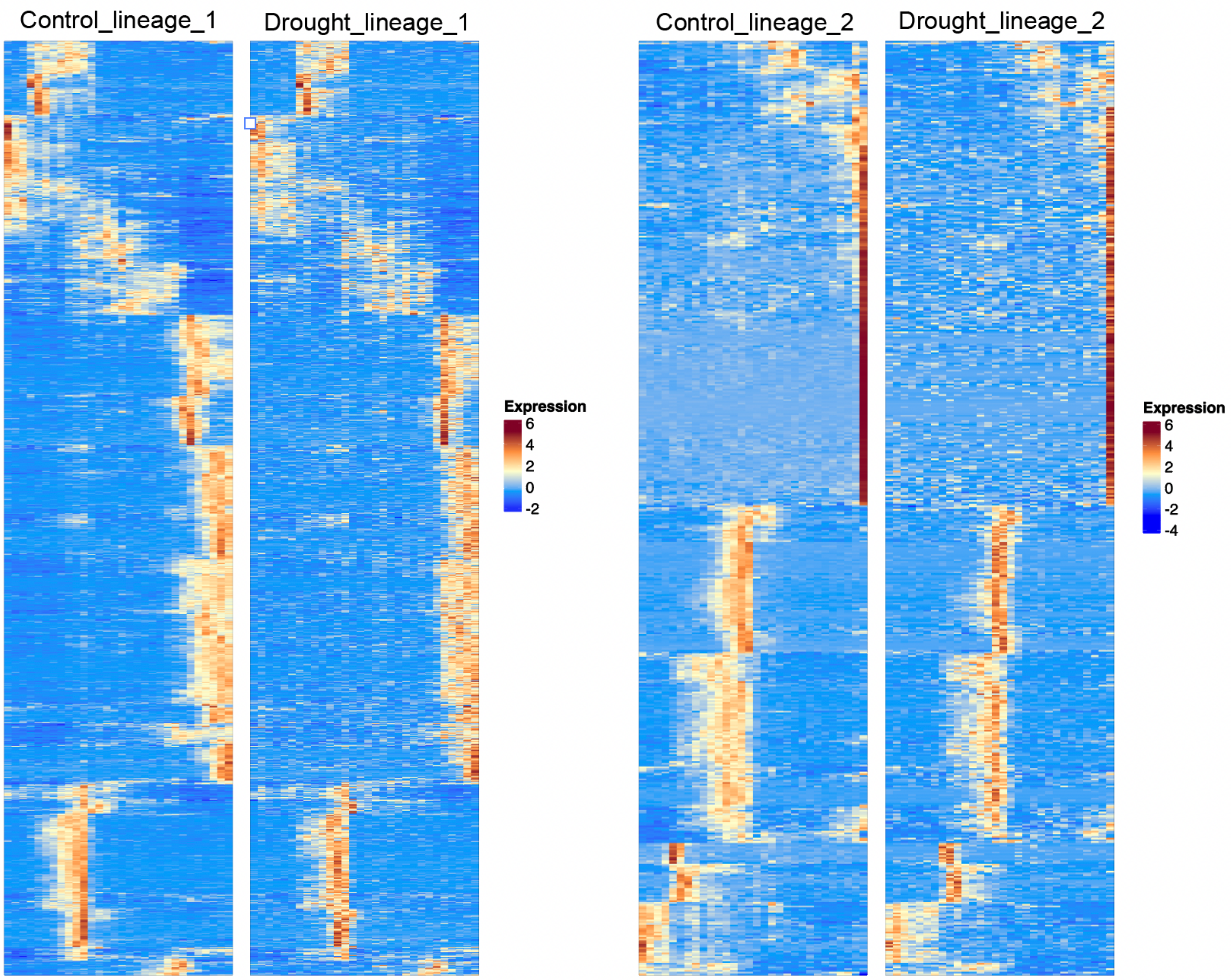


**Supplementary Fig. 7.** Heatmap showing the expression of the genes that showed significant changes along Lineage 1 (1023 genes) and Lineage 2 (537 genes) in both well-watered (control) and drought conditions (FDR < 0.05), as identified by tradeSeq using the trajectory shown in Fig. S6 as input.

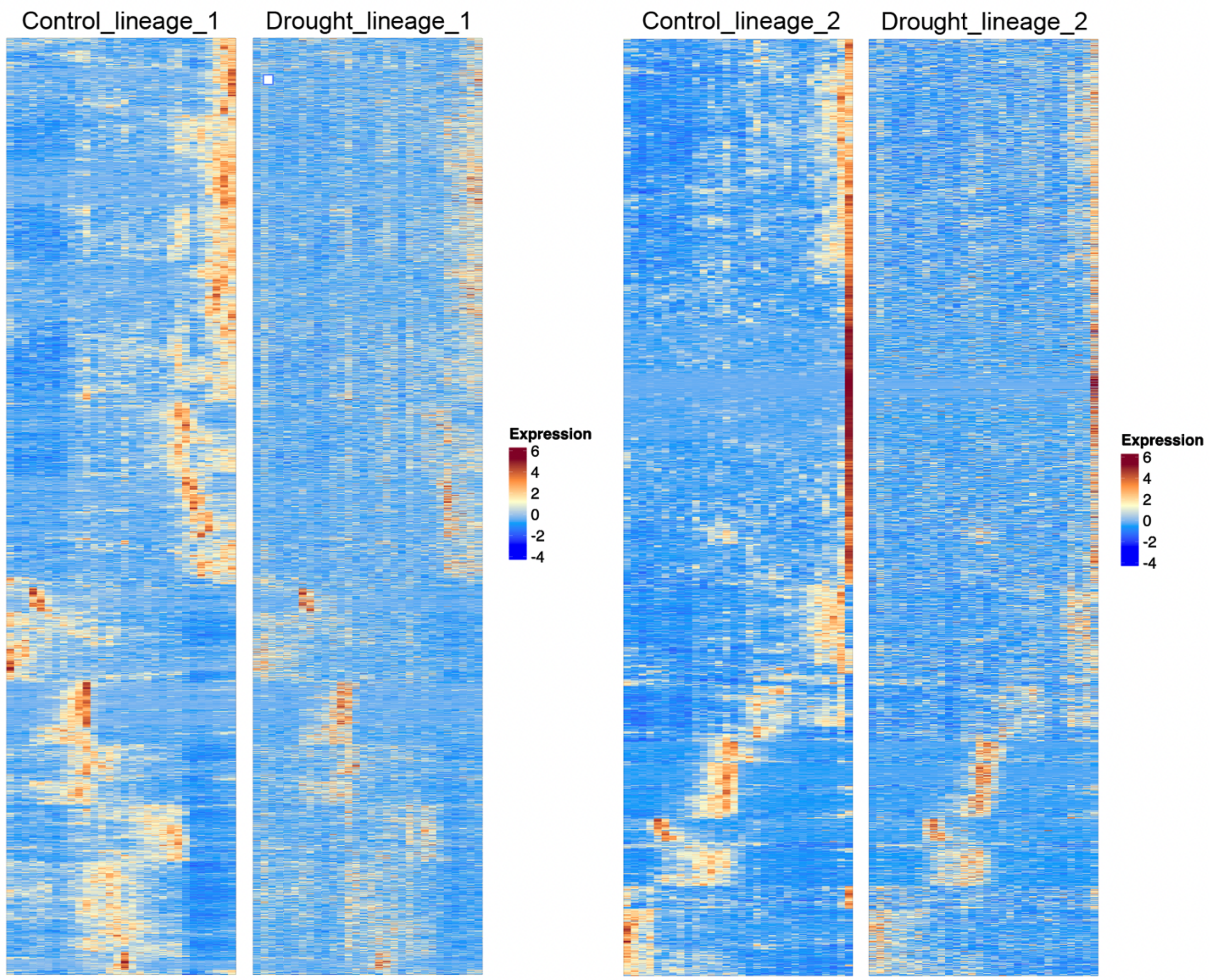


**Supplementary Fig. 8.** Heatmap showing the expression of the genes that showed significant changes along Lineage 1 (4001) and Lineage 2 (2450) only in the well-watered (control) condition and not under drought (FDR < 0.05), as identified by tradeSeq using the trajectory shown in Fig. S6 as input.

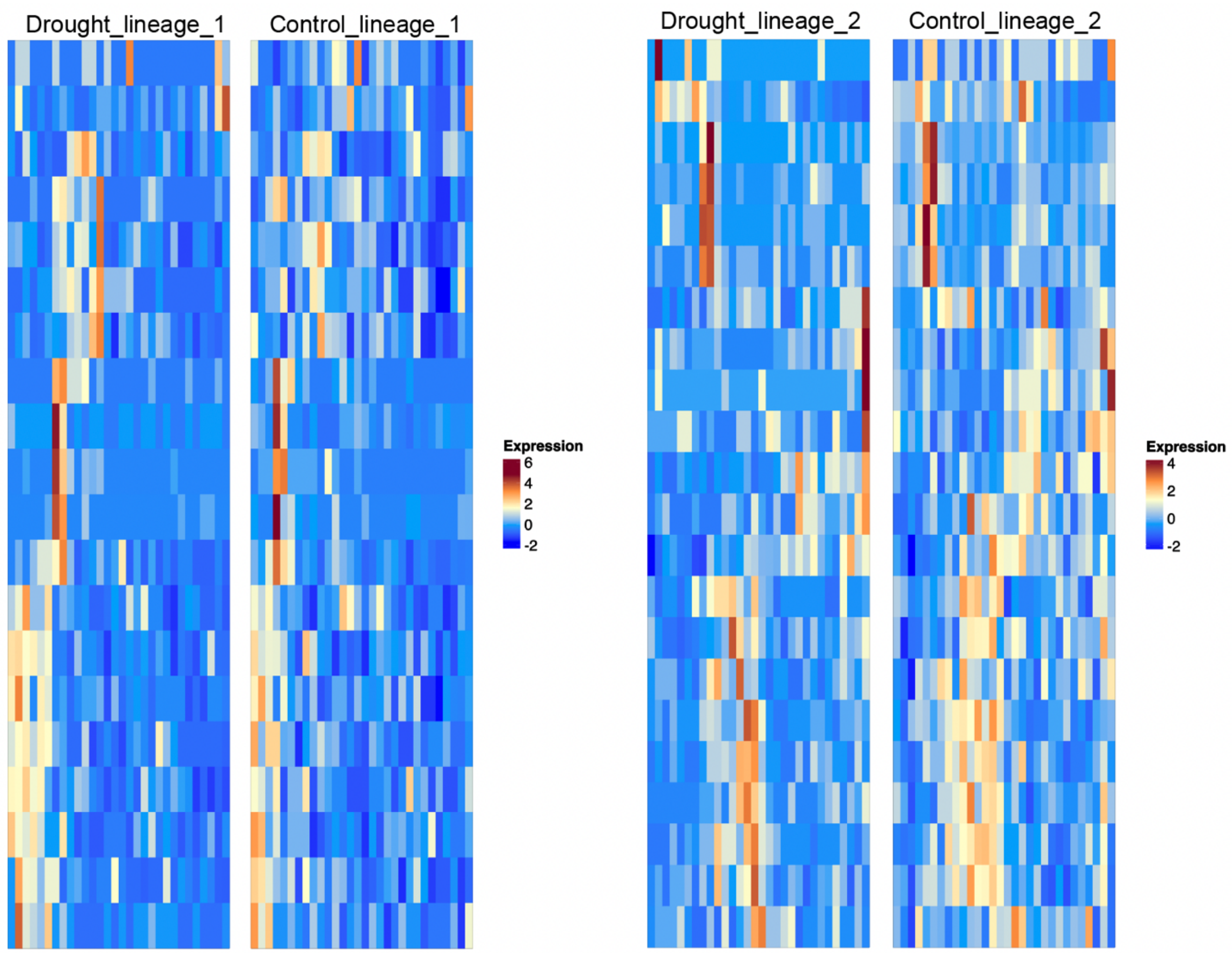


**Supplementary Fig. 9.** Heatmap showing the expression of the genes that showed significant changes along Lineage 1 (20) and Lineage 2 (22) only under drought and not in the well-watered (control) condition (FDR < 0.05), as identified by tradeSeq using the trajectory shown in Fig. S6 as input.

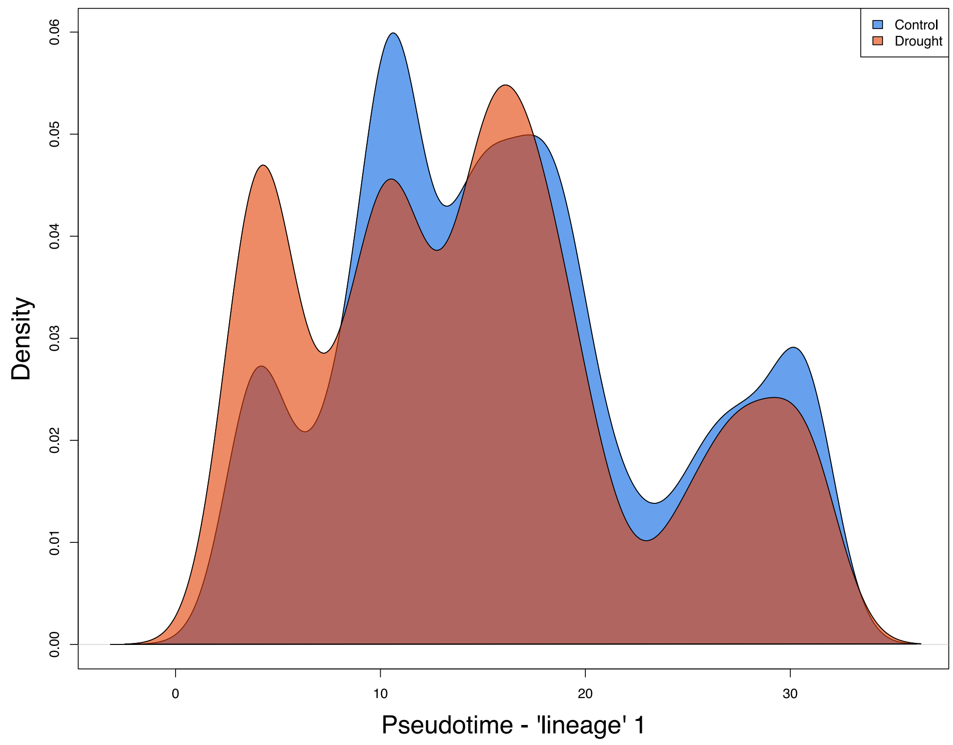


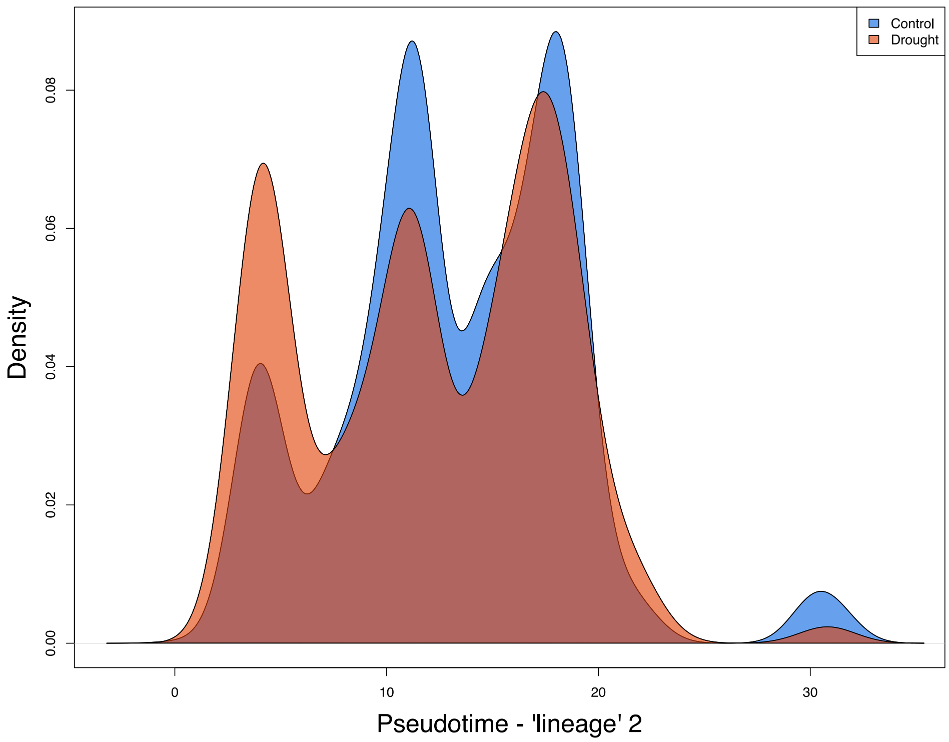


**Supplementary Fig. 10.** Density plot showing the distribution of cells along each lineage of the trajectory representing the secondary xylem (enriched in vessels) development, according to their pseudotime. In blue is the distribution of cells from the well-watered (control) condition, and in red is the distribution of cells from the drought condition. In both lineages, there is a higher concentration of cells from drought in the earlier stages of the trajectory, as shown by their lower pseudotimes values. This suggests a delay in the progression of cell differentiation under drought.

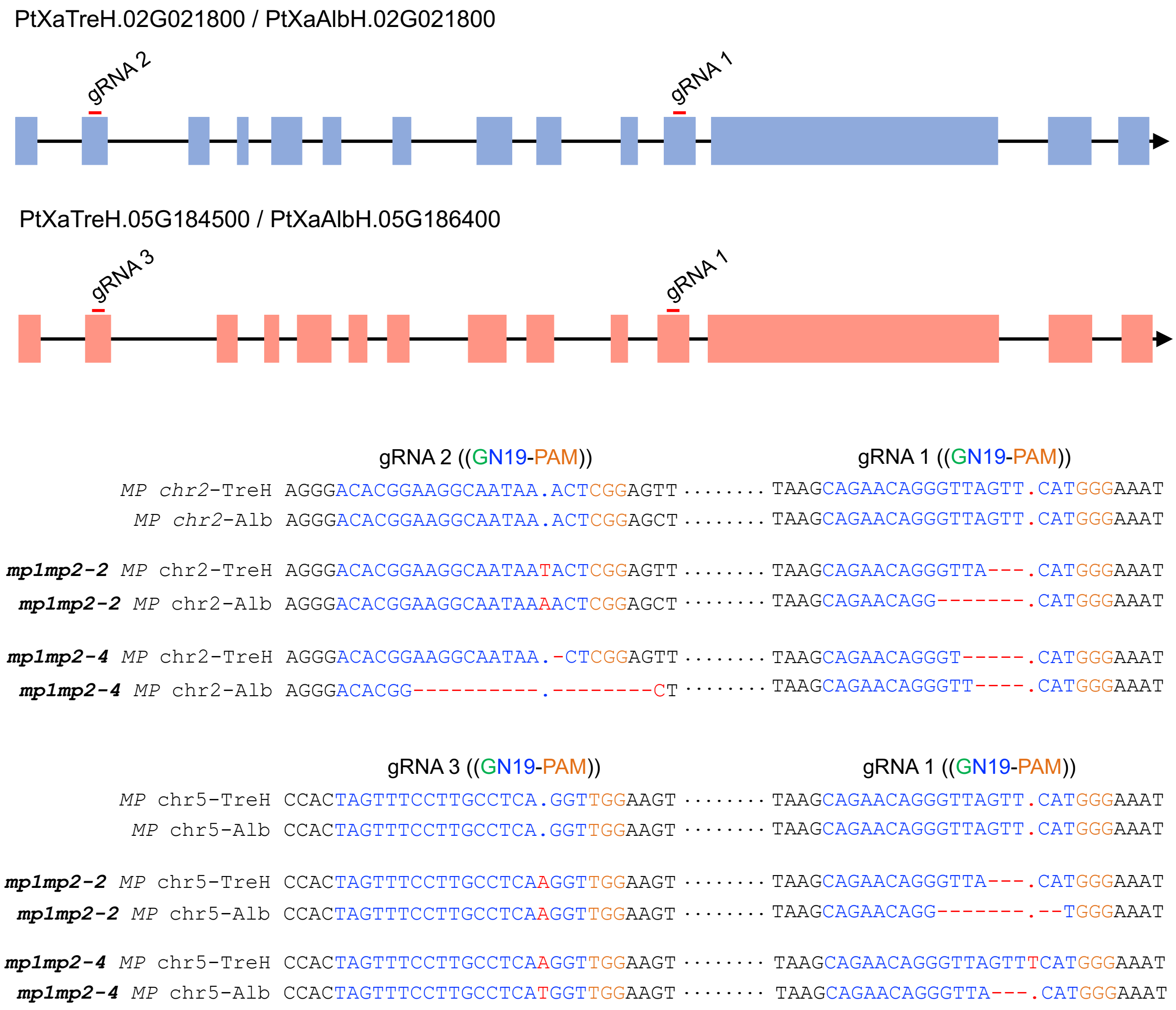


**Supplementary Fig. 11.** CRISPR/Cas9-mediated mutations of the two *MP*-like genes in *Populus tremula* × *alba* INRA clone 717 1B4. *Agrobacterium tumefaciens*-mediated transformation generated transgenic lines expressing Cas9 nuclease and three guide RNAs (gRNAs, two gRNAs targeting each *MP*-like copy). Each gRNA contains a sequence of 19 nucleotides (N19) that bind to the target DNA, guiding Cas9 to the two copies of *MP* CDS sites (exons). The protospacer adjacent motif (PAM, NGG, orange) is located at 3′ downstream of the target sequence and is necessary for the Cas9 to cleave the target site to generate a double-strand break (DSB). When a DSB was repaired, mistakes, such as small insertions and deletions, were created in or near the target locus. Two independent lines (*mp1 mp2-2* and *mp1 mp2-4*), containing mutations in both alleles of the two copies of *MP*-like genes, were selected for their functional characterization.

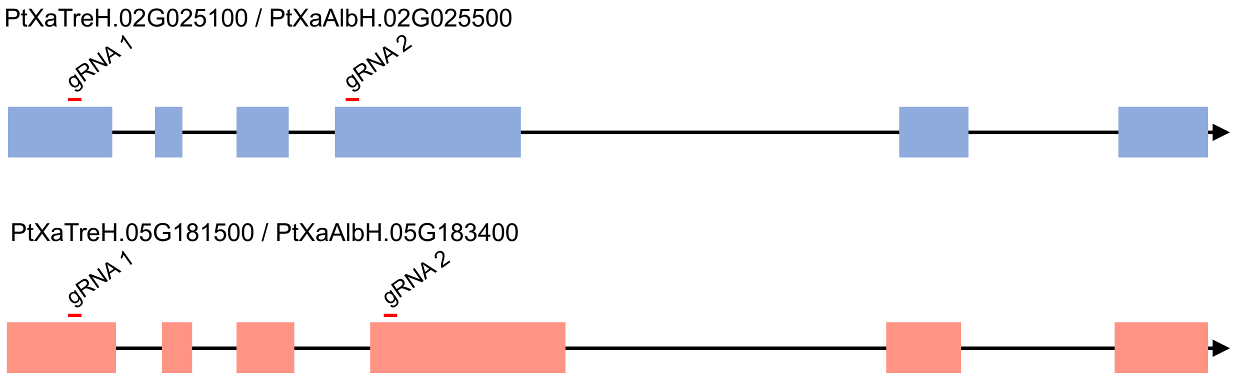


```
                              gRNA 1 ((GN19-PAM))                  gRNA 2 ((GN19-PAM))

              WAT chr2-TreH CTGCCCTAAA.TATGGGAGTTAGCAAGCTTG······AGAGATGGAATAGCAAAAGT.ACTAGGAACT
               WAT chr2-Alb CTGCCCTAAA.TATGGGAGTTAGCAAGCTTG······AGAGATGGAATAGCAAAAGT.ACTAGGAACC

wat1wat2-1 WAT chr2-TreH CTGCCCTAAA.--TGGGAGTTAGCAAGCTTG······AGAGATGGAATAGCA-----.-CTAGGAACT
 wat1wat2-1 WAT chr2-Alb CTGCCCTAAAATATGGGAGTTAGCAAGCTTG······AGAGATGGAATAGCAAA---.ACTAGGAACC

wat1wat2-9 WAT chr2-TreH CTGCCCTAAAATATGGGAGTTAGCAAGCTTG······AGAGATGGAATAGCAAAAGT.ACTAGGAACT
 wat1wat2-9 WAT chr2-Alb CTGCCCTAAAATATGGGAGTTAGCAAGCTTG······AGAGATGGAATAGCAAAAGTTACTAGGAACC

              WAT chr5-TreH CTGCCCTAAA.TATGGGAGTTAGCAAGCTTG······AAAGATGGAATAGCAAAAGT.ACTAGGAACT
               WAT chr5-Alb CTGCCCTAAA.TATGGGAGTTAGCAAGCTTG······AAAGATGGAATAGCAAAAGT.ACTAGGAACT

wat1wat2-1 WAT chr5-TreH CTGCCCTAAA.TATGGGAGTTAGCAAGCTTG······AAAGATGGAATAGCAAAA--.ACTAGGAACT
 wat1wat2-1 WAT chr5-Alb CTGCCCTAAA.TATGGGAGTTAGCAAGCTTG······AAAGATGGAATAGCAAAAGT.-CTAGGAACT

wat1wat2-9 WAT chr5-TreH CTGCCCTAAA.      INSERTION 668 BASE PAIRS **        T.ACTAGGAACT
 wat1wat2-9 WAT chr5-Alb CTGCCCTAAA.TATGGGAGTTAGCAAGCTTG······AAAGATGGAATAGCAAAAGTTACTAGGAACT
** INSERTION 668 BASE PAIRS
ACTTTTGCTATTCCATCTTTTCTGTTTATCCTCACTTTCTCTATCCTGCATCATCAAAGAAAATAACACGTTTCAGAAAACGAAAATTAGGTGTAAAA
CGATGCATTTACACGCTCCAATCATATGTATGTGCATAAAATTAAACGTGTGATGCAATAAGAGTCTAAAAGAGAGAGAAAGGGAGAGGAGAGAGGGA
GAGGGAGGGGGCTTGCCGGAGTAGAGCTGCCATGAGAAAGGTAATAGCAGGGACAGAGTTTTGGATTGCTGATGCGAAGGTGGGTGAAGTGTTCTCCA
GACCCAGCAAGTAGAATCCTTGGTTTGCTGTTATTCTGTTCACATGGGCAACAAGAAGTGAAAATGAGCGGTCATTGTTTTTTATTTTTATTTTCATT
TCAAATTATTTGATATATTTTGGGTCATACCCAACAAGTGCAAGGAAGAAGAACTGAAGGACAAAGTTGAGAGTGAGCGCTGGCCTCTCCTTCCTGAT
CGAACAAAGAGAAAAGAATAAATACTCCTTGAAAATATATAAAGGATGACTGAAGATAAAATGAAAGGGTTAGATTAATGGATAGCAGTACTTTTCCA
GAAAATATGCAAAGGGAAGGAGCAGAAGCAAAGCGATGATGTTCCTATATACTGGGAAAACAAGCTTGCTAACTCCCATA
```

**Supplementary Fig. 12.** CRISPR/Cas9-mediated mutations of the two *WAT*-like genes in *Populus tremula* × *alba* INRA clone 717 1B4. Agrobacterium tumefaciens-mediated transformation generated transgenic lines expressing Cas9 nuclease and two guide RNAs (gRNAs). Each gRNA contains a sequence of 19 nucleotides (N19) that bind to the target DNA, guiding Cas9 to the two copies of MP CDS sites (exons). The protospacer adjacent motif (PAM, NGG, orange) is located at 3′ downstream of the target sequence and is necessary for the Cas9 to cleave the target site to generate a double-strand break (DSB). When a DSB was repaired, mistakes, such as small insertions and deletions, were created in or near the target locus. Two independent lines (*wat1 wat2-2* and *wat1 wat2-4*) containing mutations in both alleles of the two copies of *WAT*-like genes were selected for their functional characterization.

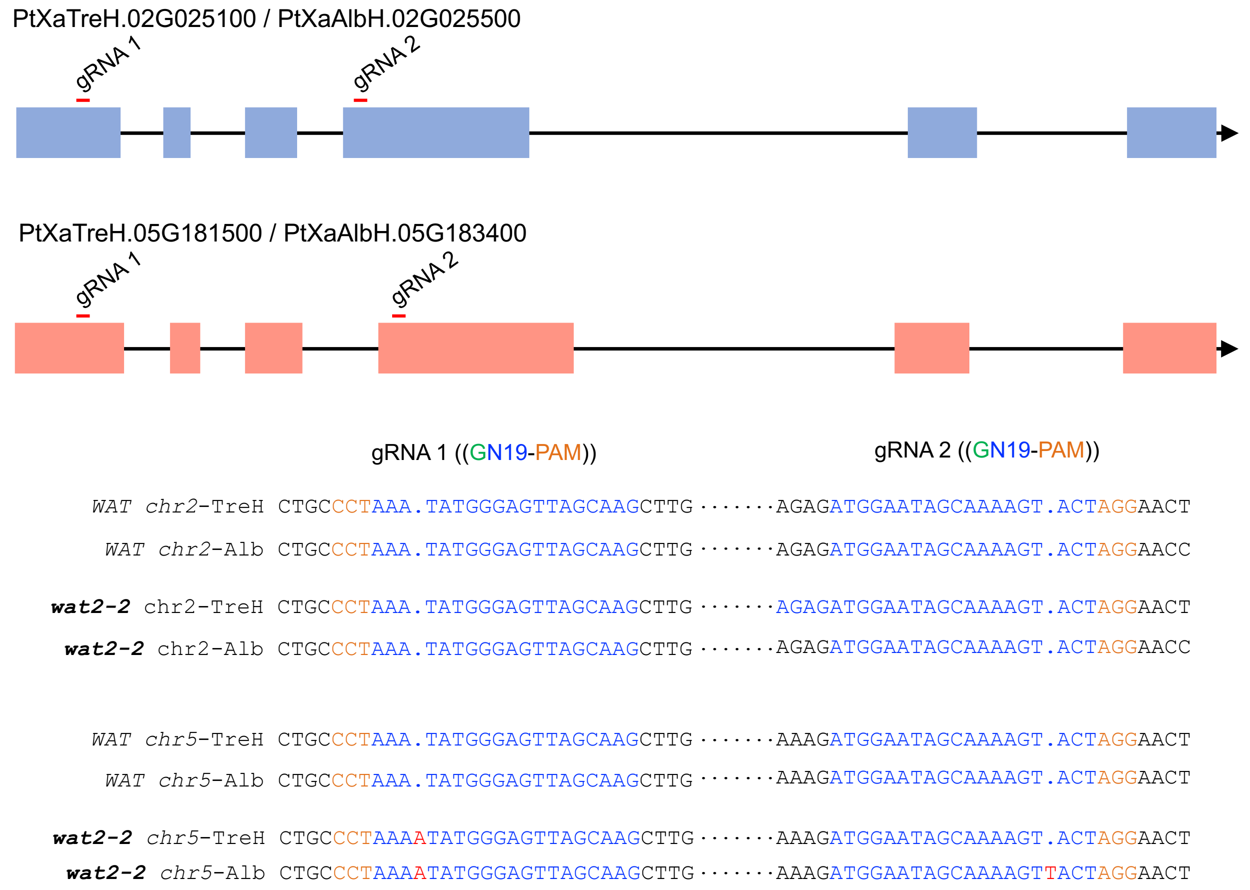


**Supplementary Fig. 13.** Sequence of the *wat2* single mutant (*wat2-2*) identified in the screening performed to identify CRISPR/Cas9-mediated mutants for *WAT*-like genes (Supplementary Fig. 12).
